# ETEC heat-labile toxin promotes β-catenin stabilization and transcriptional reprogramming to disrupt intestinal epithelial differentiation

**DOI:** 10.1101/2025.10.02.680135

**Authors:** Alaullah Sheikh, Bipul Setu, Nikhilesh Joardar, John C. Martin, Bruce A. Rosa, Makedonka Mitreva, James M. Fleckenstein

## Abstract

Enterotoxigenic *Escherichia coli* (ETEC), defined by their production of heat-labile (LT) and heat-stable (ST) enterotoxins, are a common cause of acute diarrheal illness in children from low- and middle-income countries, and are also linked to long-term sequelae such as malnutrition, and growth impairment. While the mechanisms underlying toxin-mediated acute diarrhea are known, the molecular events involved in ETEC related sequelae remain unclear. Here, we demonstrate that the ETEC heat-labile toxin (LT) profoundly remodels intestinal epithelial composition and function through modulation of WNT/β-catenin signaling. Using human ileal enteroids, we demonstrate that LT stabilizes β-catenin independently of WNT ligands, promotes its nuclear accumulation, and enhances TCF/LEF-driven transcription. Single-cell transcriptomic analyses reveal that LT increases intestinal proliferation by enhancing cell cycle activity across all epithelial lineages, thus disrupting epithelial composition by expanding proliferative progenitor populations at the expense of absorptive enterocytes. Simultaneously, LT impairs epithelial maturation and suppresses transcriptional programs required for nutrient absorption and differentiation. Together, these findings identify LT as a potent driver of intestinal epithelial reprogramming, providing mechanistic insight into how ETEC infection may drive long-term consequences beyond acute diarrhea and may inform strategies to prevent major sequelae, including malnutrition, that affect millions of children worldwide.

## Introduction

The enterotoxigenic *E. coli* (ETEC) are a diverse group of diarrheal pathogens defined by their production of heat-labile (LT) and/or heat-stable (ST) enterotoxins^1^. The ETEC pathovar is thought to be responsible for hundreds of millions of cases of diarrheal illness each year, primarily among young children of low-middle income countries (LMICs)^2^. Children of LMICs frequently suffer multiple episodes of ETEC diarrhea before in the first few years of life^3^, and ETEC are a leading cause of death due to diarrheal illness in children under the age of five years^4^. While the death rate from acute diarrheal illness has declined with advances in oral rehydration therapy and other measures, children with ETEC diarrhea are also at increased risk of death after resolution of the initial episode^5^. Moreover, children with moderate-severe diarrhea^6^ as well as less severe diarrhea^7^ due to ETEC inexplicably suffer long-term sequelae including from a condition referred to as environmental enteric dysfunction (EED) characterized histologically by atrophy of small intestinal villi accompanied by expansion of crypts^8,9^, and clinically by poor growth and malnutrition. Unfortunately, in the absence of a licensed vaccine, morbidity due to acute diarrheal illness as well as long-term sequelae of impaired growth and malnutrition have continued unabated^7^.

The fundamental molecular pathogenesis of acute diarrheal illness caused by ETEC is well-established. Delivery of LT and ST activates production of cAMP and cGMP cyclic nucleotides, respectively. Increases in intracellular cAMP leads to activation of protein kinase A (PKA) while cGMP activates protein kinase G (PKG). Kinase-mediated phosphorylation of cellular ion channels in enterocyte membranes leads to the net export of salt and water into the intestinal lumen, culminating in profuse watery diarrhea characteristic of ETEC.

In contrast, the sequelae linked to ETEC are still poorly understood. Recently, however we demonstrated that heat-labile toxin modulates the transcription of hundreds of genes in target intestinal epithelial cells, including those responsible for the biogenesis of enterocyte microvilli responsible nutrient absorption^10^. Moreover, repeated challenge of infant mice with toxigenic ETEC leads to impaired growth^10^ similar to observations in children. These studies provided the first suggestion that ETEC toxins might drive molecular events that culminate in sequelae.

Nutrient absorption in the small intestine depends on tightly regulated epithelial differentiation and maturation processes that generate functionally specialized cell types. Intestinal stem cells (ISCs), residing at the base of crypts, undergo proliferation and lineage specification to give rise to absorptive enterocytes and secretory lineages including goblet, Paneth, and enteroendocrine cells^11^. As these cells migrate along the crypt-villus axis, they progressively differentiate, acquiring mature phenotypes and functional attributes^12^. In particular, enterocytes undergo a stepwise transcriptional and morphological maturation program culminating in the expression of solute carrier (SLC) transporters, and other molecules essential for nutrient uptake and barrier function^13^. Theoretically, disruption of this developmental trajectory would compromise nutrient absorption and contribute to malnutrition among children from low-resource settings^8,14-17^.

Intestinal epithelial maturation is tightly regulated. The luminal surface of the small intestine is comprised of millions of villi, tiny (∼0.5 mm long) fingerlike projections that are surrounded at their base by invaginations known as crypts. As epithelial cells covering the villi are propagated from intestinal stem cells (ISC) residing in the crypts they quickly differentiate into mature epithelial cells, with absorptive enterocytes covered in microvilli making up the majority. These mature cells on the villous surface undergo continual renewal over a period of 3-5 days^18^ as they are constantly replenished by the population of crypt stem cells. Differentiation of ISC into enterocytes and other cell types along the crypt-villus axis is controlled by a number of signaling molecules including WNT^19-21^.

High concentrations of WNT generated by adjacent Paneth cells^22^ favor the maintenance and proliferation of crypt (ISC) populations. In the absence of WNT, a “destruction complex” comprised of GSK-3 β, CK1α, APC, and Axin2 engages β-catenin. GSK-3 β phosphorylation of β-catenin targets it for ubiquitination^23^ and removal from the cytoplasm. Binding of WNT to its cognate LRP5/6-Frizzled co-receptors on the cell surface activates Dishevelled, a cytoplasmic protein that antagonizes the destruction complex^24^. This permits accumulation of monomeric β-catenin in the cytoplasm^25^ and ultimately its translocation to the nucleus where it binds to T-cell factor (TCF) and lymphoid enhancer factor (LEF) transcription factors^26^, leading to activation of WNT/TCF target genes responsible for maintenance of the crypt progenitor cells. In addition to canonical WNT signaling, β-catenin can be stabilized both directly and indirectly by modulating features of the destruction complex. Here, we demonstrate that heat-labile enterotoxin of ETEC enhances β-catenin stabilization to activate downstream signaling in small intestinal epithelia, disrupting differentiation and maturation required for efficient nutrient uptake.

## Results

### Heat-labile toxin activates transcription of multiple WNT-responsive genes

Bioinformatic analysis (CompBio v2.9) of earlier bulk RNAseq data^10^ from small intestinal enteroids treated heat-labile toxin revealed that transcription of genes governed by the WNT-β-catenin pathway were significantly upregulated in response to LT (figure 1a). Upregulated genes included Axin2, RNF43, MYC, SOX9, cyclin D1 (CCND1), EphB2, LGR5, PROM1, DKK1, CLDN1, TSPAN8, along with a number of other genes governing the intestinal cell differentiation including OLFM4, CD24, REG4, XBP1, and ATOH1^21,27,28^. Using CEACAM6 as a positive control^29^ for LT-induced gene expression, we validated impact on genes in this pathway by qRT-PCR (Figure 1b). In keeping with these results, we also observed cellular accumulation of β-catenin following treatment of polarized small intestinal enteroids with LT (Figure 1c,d).

**Figure 1.**
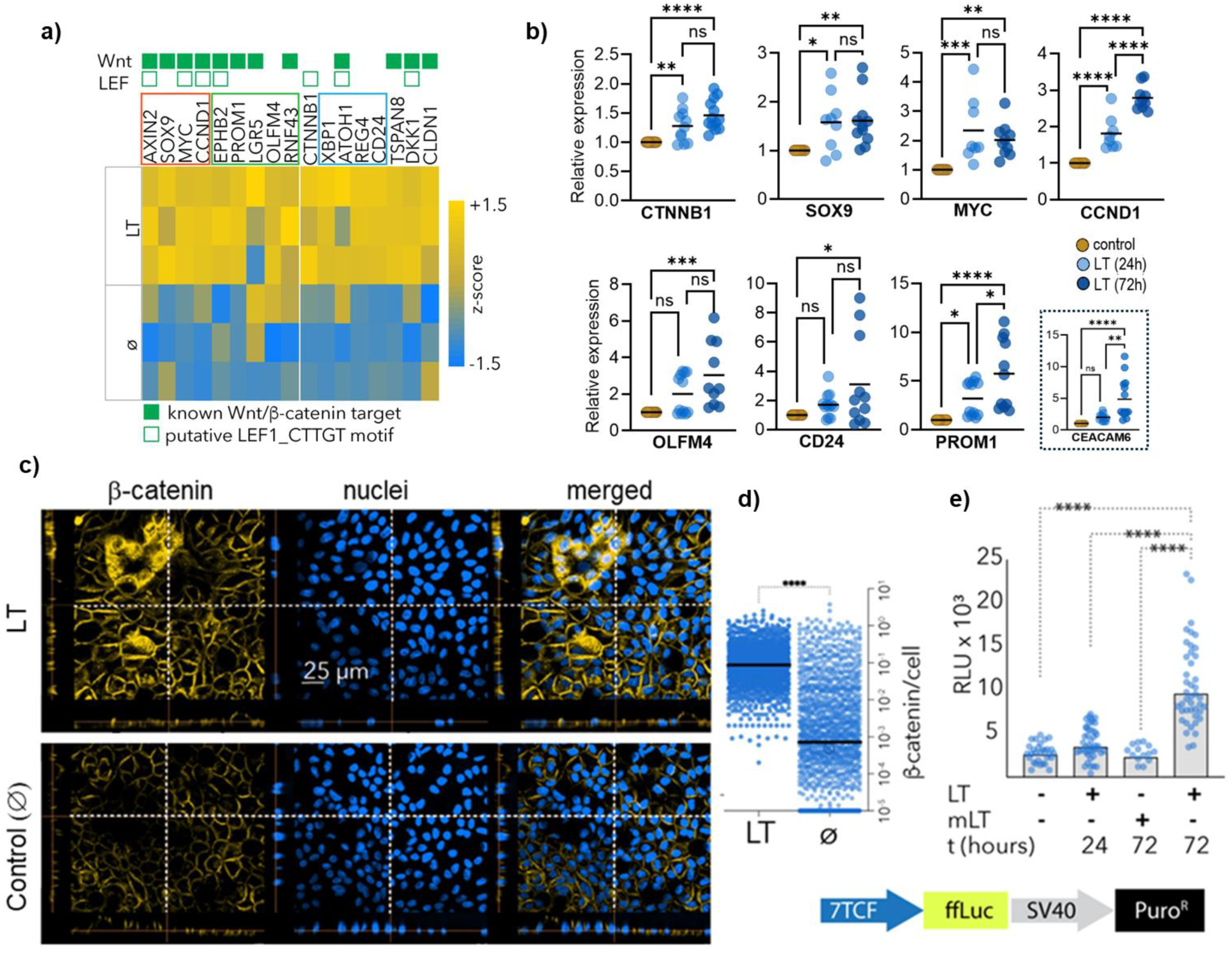
LT activates WNT/β-catenin signaling in human intestinal epithelial cells. **a**) Heatmap showing differential expression (z-score) of WNT/β-catenin signaling components and target genes in control (Ø) and LT-treated enteroids based on bulk RNA-seq. Closed green symbols signify genes known to be modulated by canonical WNT-β-catenin pathway as profiled (https://web.stanford.edu/group/nusselab/cgi-bin/wnt/target_genes). Open green symbols indicate genes possessing an upstream 3’UTR CTTTGT motif for the LEF1: lymphoid enhancer-binding factor http://www2.stat.duke.edu/∼sayan/genesets/Jan2006/cards/C3/CTTTGT_V_LEF1_Q2.html. **b**) RT-qPCR validation of selected genes showing significantly increased expression following LT treatment. Gene expression was measured at 24 and 72 hours post-treatment. Data are presented as expression levels normalized to GAPDH and shown relative to the untreated control. Inset: CEACAM6 expression is included as a positive control for LT-mediated induction. Statistical comparisons were made using Kruskal-Wallis; ns = not significant, *p < 0.05, **p < 0.01, ***p < 0.001, ****p < 0.0001. **c**) Representative confocal immunofluorescence images of β-catenin (yellow) and nuclei (blue) in LT-treated and control enteroids showing increased nuclear β-catenin accumulation in LT-treated samples. Scale bar, 25 μm. **d**) Quantification of nuclear β-catenin intensity per cell from images shown in panel (c). Each dot represents an individual nucleus. ****p<0.0001 Mann-Whitney (two-tailed). **e**) Luciferase reporter activity (in Relative light units, RLU) following treatment of Hu235D::7TFP cells with heat-labile toxin (LT) or mutant, catalytically inactive toxin (mLT, E112K). The data shown are combined from 3-7 independent experiments. Each dot represents signal per well and each Bar represents geometric mean. Statistical analysis by one-way ANOVA (Kruskal-Wallis), ****p<0.0001.

To functionally assess transcriptional modulation by β-catenin, we stably transfected small intestinal enteroids with a lentivirus vector construct (7TFP)^30^ which contains 7 copies of the TCF/LEF transcription factor DNA-binding site 5’-CCTTTGATC-3’ upstream of firefly luciferase^31^. Treatment of 7TFP carrying reporter enteroids with LT resulted in significant increases in TCF/LEF-mediated transcriptional activity (Figure 1e). These effects were abrogated when the enzymatically inactive mutant form of LT (mLT) was used, indicating that catalytic activity of LT is required for this activation.

### Heat-labile toxin stabilizes cytosolic β-catenin to facilitate nuclear translocation

In canonical WNT signaling pathways, β-catenin accumulation in the nucleus permits engagement of TCF/LEF transcription factors to activate transcription of downstream genes (Figure 2a). Interestingly, activation of cyclic nucleotide pathways has the potential to interfere with degradation of β-catenin. PKA phosphorylation of β-catenin at position Ser_675_ inhibits β-catenin ubiquitination, thereby permitting its accumulation^32^, while activation of PKA can also result in phosphorylation of GSK-3β at Ser_9_ resulting in its inhibition^33-37^. Similarly, cGMP activation of PKG also results in phosphorylation of GSK-3β at Ser^38,39^ to inhibit enzymatic activity. Accordingly, we observed time-dependent increases in total β-catenin in both cytoplasmic and nuclear compartments (Figure 2b-e). In parallel, LT induced phosphorylation of β-catenin at Ser_675_ (Figure 2b-e, Supplementary figure 1), a known activating modification that promotes β-catenin nuclear localization and transcriptional activity^40^, further supporting functional stabilization.

**Figure 2.**
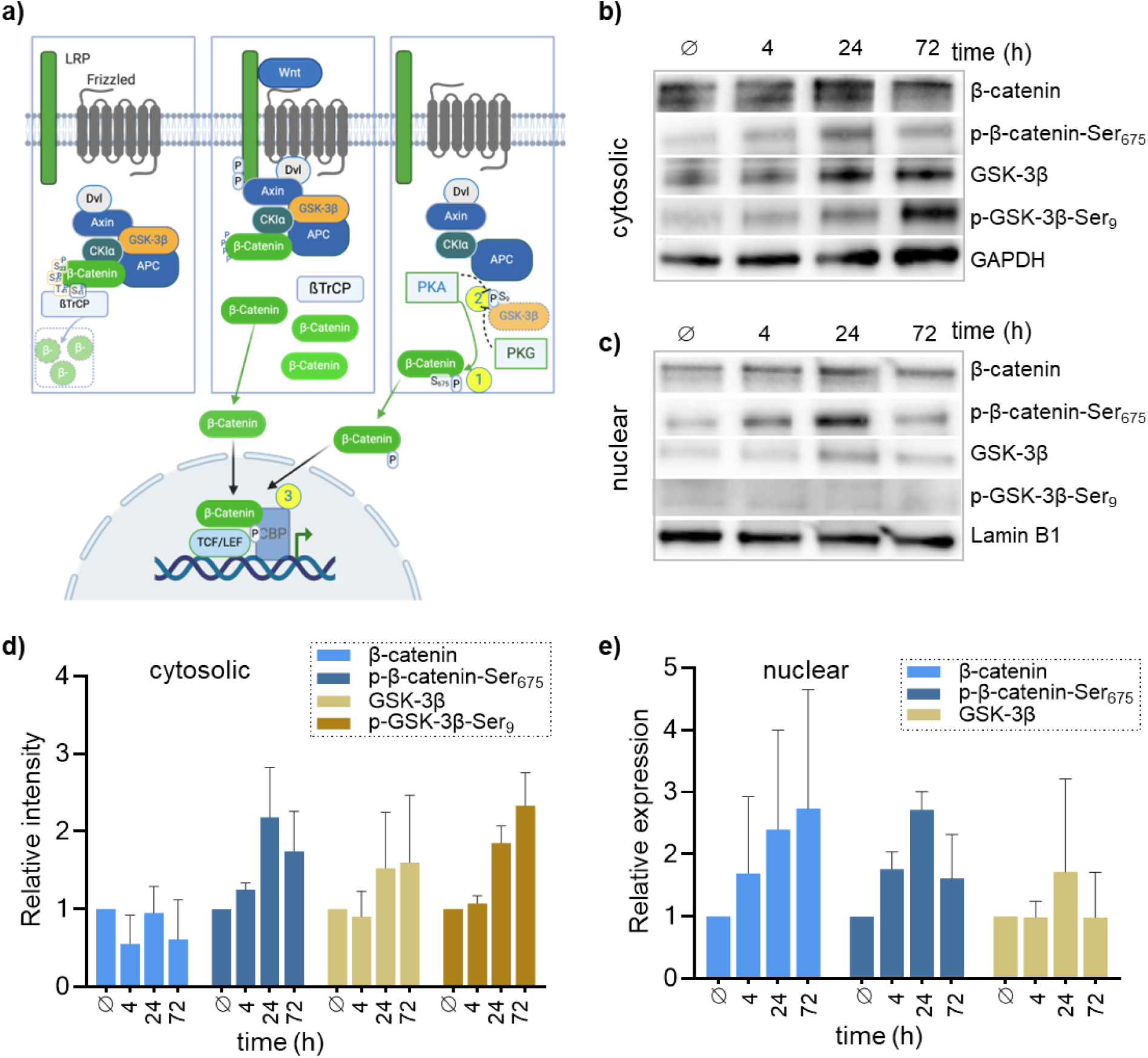
Heat-labile toxin stabilizes β-catenin signaling independent of WNT. **a)** Diagram illustrating canonical WNT–β-catenin signaling (left and middle panels). Panel at right depicts anticipated impacts of PKA-mediated, WNT-independent signaling. **b**) Representative immunoblots of cytoplasmic and **c**) nuclear fractions from LT-treated enteroids at indicated time points (4, 24, 72 hours) or untreated controls (Ø). Blots were probed for total β-catenin, phospho-β-catenin (Ser_675_), total GSK3β, and phospho-GSK3β (Ser_9_). GAPDH and lamin B1 served as loading controls for cytoplasmic and nuclear fractions, respectively. **d-e)** Graphs represent densitometry measurements of immunoblots of cytoplasmic (**d**) and nuclear (**e**) fractions, normalized to respective loading controls, summarizing 4 independent experiments. Bars represent geometric means ± SD.

In addition, we found that LT modulated phosphorylation of GSK-3β, a central negative regulator of β-catenin stability. Although total protein levels remained relatively unchanged, GSK-3β phosphorylation at the Ser_9_ residue increased significantly in LT-treated samples over time (Figure 2b, d).

Collectively, these data demonstrate that heat-labile toxin acts to stabilize β-catenin independent of WNT by directed post translational modification of β-catenin and elements of the destruction complex^32,41^. Interestingly, we also noted that transcription of the *CTNNB1* gene encoding β-catenin was itself upregulated in response to LT, potentially related to the presence of cAMP response elements^42^ in the CTNNB1 promoter^43^.

### Heat-labile toxin disrupts epithelial cell composition

Given the central role of WNT/β-catenin signaling in regulating intestinal stem cells^20^ and their differentiation to different epithelial cell types^18^, we next investigated whether LT-induced stabilization of β-catenin impacts epithelial cell dynamics. In the intestine, proliferative expansion without coordinated differentiation can shift the cellular composition and impair epithelial function^44^. To investigate whether heat-labile toxin perturbs this balance, we used single-cell RNA sequencing to examine the impact of LT on differentiation of human small intestinal enteroids. Control (Ø) and LT-treated enteroid cells were harvested and subjected to Illumina NovaSeq X Plus single-cell RNA sequencing, followed by analysis with Seurat (v5.0)^45^. After quality control filtering, 1,963 cells represented by an average of 14,342 reads and 3,266 genes were sequenced in the control sample, and 2207 cells, an average of 14,099 reads and 3283 genes in the LT sample. To assess changes in epithelial cell composition, we then performed unsupervised graph-based clustering of single-cell transcriptomes from control and LT-treated enteroids (resolution setting of 0.92, defining 14 clusters), followed by annotation using canonical lineage markers^11,13^, confidently identifying all expected major cell types (Figure 3a; Table 1; **Supplementary dataset 1**).

**Figure 3.**
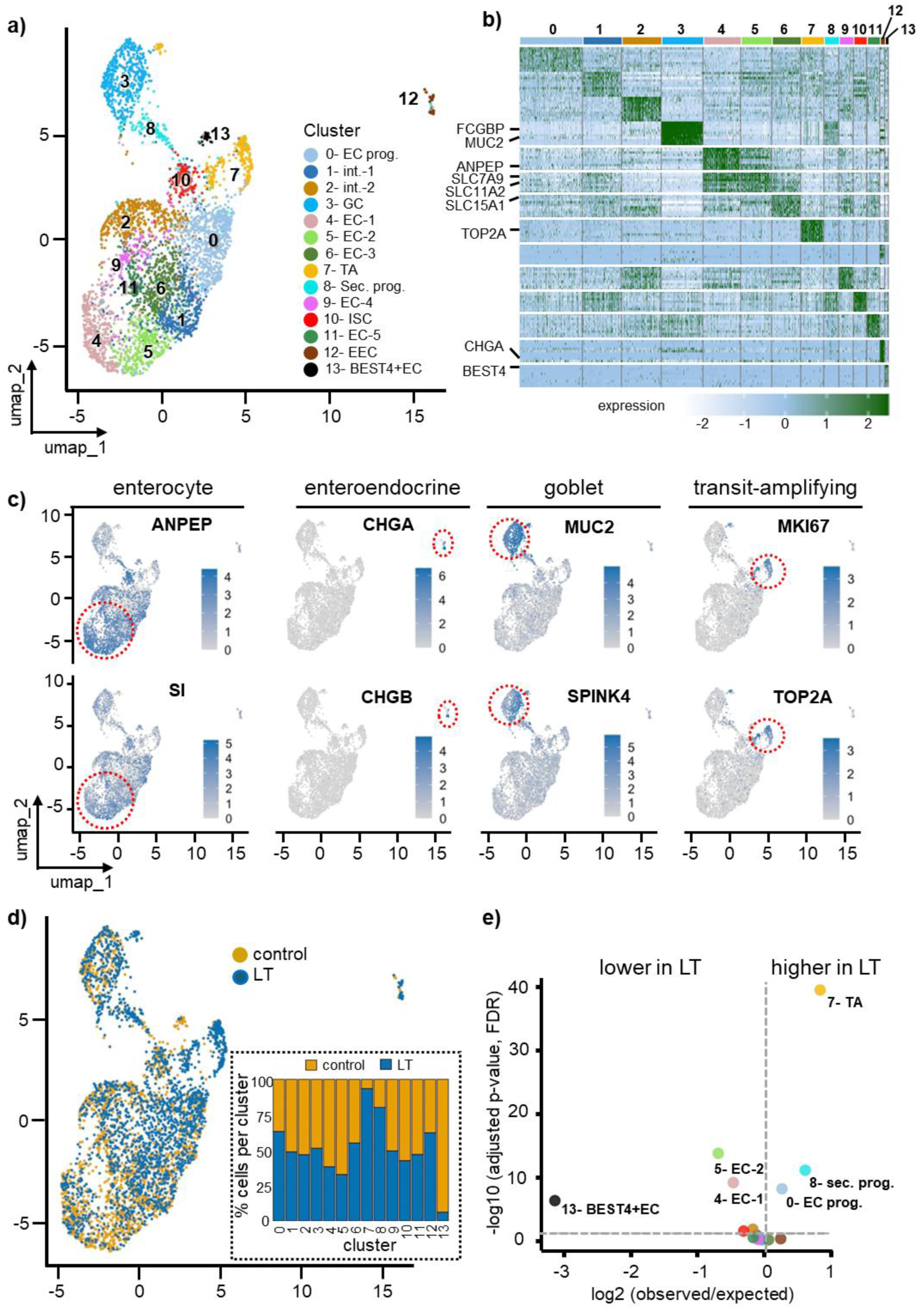
LT treatment reshapes intestinal epithelial cell composition. **a**) UMAP projection of single-cell transcriptomes from control and LT-treated ileal enteroids, colored by cluster identity. Unsupervised clustering identified 14 transcriptionally distinct epithelial populations, which were annotated based on lineage marker gene expression. These include enterocyte progenitors, intermediate cells, secretory lineages (goblet, enteroendocrine, and Paneth-like), stem cells, and multiple stages of differentiated enterocytes. **b**) Heatmap showing the relative expression levels of the top 10 marker genes per cluster, showing distinct transcriptional signatures corresponding to absorptive, secretory, intermediate, and proliferative compartments. Expression values represent SCTransform-normalized Pearson residuals. **c**) UMAP feature plots showing expression of known cell type specific markers. Enterocyte markers: ANPEP, SI, ALPI; goblet cell markers: MUC2, SPINK4, TFF3; enteroendocrine cell markers: CHGA, CHGB, THP1; transit-amplifying cell markers: MKI67, TOP2A, PCNA. Red dotted circles highlight specific cell clusters with high expression values. **d**) UMAP showing the distribution of cells from control (orange) and LT-treated (blue) samples, revealing a shift in cell type representation. Inset: Stacked bar plot showing the proportion of cells from each sample (control vs LT) within each cluster. **e**) Differential abundance analysis for epithelial cell clusters in LT-treated versus control enteroids. Each point represents an epithelial cell cluster. The x-axis shows the log₂ ratio of observed to expected cell counts in LT-treated enteroids, and the y-axis shows the –log₁₀(FDR-adjusted p-value). Vertical and horizontal dashed lines indicate the direction of change (assigned based on whether the observed LT count exceeded or fell below the expected value) and statistical significance thresholds, respectively. Clusters to the right of the vertical line are enriched in LT-treated samples, whereas clusters to the left are reduced. Significantly altered clusters are labeled. Statistical analysis was performed per cluster using a Chi-square test with FDR correction (significant if FDR <0.05). BEST4+EC, BEST4 positive enterocyte; EC, enterocyte; EC prog.- enterocyte progenitor; EEC, enteroendocrine cell; GC, goblet cell; ISC, intestinal stem cell; int., intermediate; sec. prog., secretory progenitor; TA, transit amplifying.

**Table 1.**
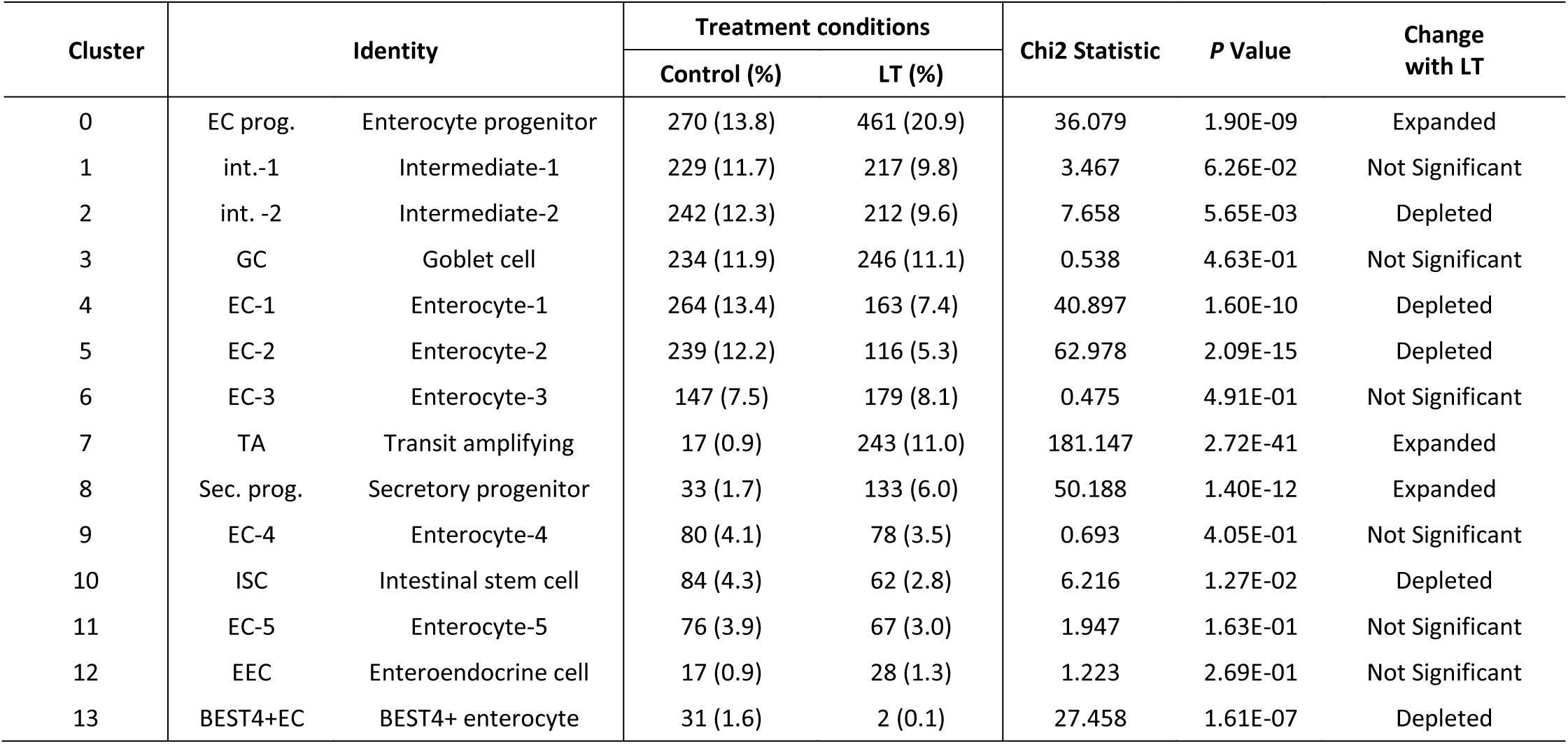
Transcriptionally-defined epithelial cell cluster compositions in ileal enteroids.

LT treatment induced striking alterations in enteroid cell composition. All major intestinal epithelial cell types, including absorptive enterocytes, goblet cells, Paneth cells, and enteroendocrine cells, along with intermediate populations likely representing transitional states along the differentiation axis, were identified in both control and LT-treated samples (Figure 3a–c). However, a highly proliferative cluster, largely absent in controls, was markedly expanded following LT exposure, while a differentiated population present only in control samples was lost (Figure 3d–e). Differential abundance analysis confirmed enrichment of early-stage progenitor and transit-amplifying populations (Clusters 0, 7, and 8) in LT-treated samples, alongside significant depletion of mature enterocytes (Clusters 4 and 5) and BEST4⁺ absorptive cells (Cluster 13) (Figure 3e; Table 1).

To further explore the identity of these transcriptionally distinct or depleted populations (Clusters 7, and 13, respectively) in LT-treated enteroids, we performed over-representation analysis (ORA). ORA of cluster 7 revealed strong enrichment for mitotic cell cycle pathways, including G2/M transition, spindle checkpoint, and DNA replication, indicating a highly proliferative state (Supplementary figure 2a). Consistently, transcription factor analysis identified dominant enrichment of E2F family regulators (E2F1, E2F2, E2F3) and their partners DP1/DP2, which are key drivers of S-phase entry and DNA synthesis (Supplementary figure 2b). Additional enriched motifs included MYC, NFY, and YY1, supporting enhanced biosynthetic and metabolic activity consistent with transit-amplifying cells. In contrast, Cluster 13 was enriched for pathways associated with terminal epithelial differentiation and absorptive function. These included mitochondrial respiration, oxidative phosphorylation, the TCA cycle, and lipid metabolism (Supplementary figure 2a). The absence of proliferative transcription factor signatures in Cluster 13 further supports its identity as a mature enterocyte population (Supplementary figure 2b). Together, these findings reveal that LT exposure promotes the expansion of transcriptionally active proliferating cells while depleting functionally mature absorptive cells, disrupting the balance between proliferation and differentiation in the intestinal epithelium.

To gain deeper insight into the transcriptional alterations underlying these shifts, we conducted differential gene expression analysis on all cell clusters, identifying thousands of differentially expressed genes (DEGs) significantly upregulated or downregulated in LT-treated enteroids compared to control (Table 2, **Supplementary dataset 2**). To assess functional implications, we performed over-representation analysis (ORA) of differentially expressed genes. Upregulated DEGs were enriched for pathways in RNA metabolism, cell cycle regulation, and stress response, consistent with a proliferative epithelial state (Supplementary figure 3). Downregulated DEGs were enriched for bile acid transport, lipid metabolism, cholesterol biosynthesis, and other absorptive functions, indicating impaired maturation and loss of differentiation (Supplementary figure 3). Gene ontology independent analysis (CompBio v2.9) likewise highlighted suppression of metabolic and transport functions, including intestinal absorption, lipid and bile metabolism, fatty acid uptake, amino acid metabolism, and cholesterol processing (Supplementary figure 4, also see **Supplementary dataset 2**). Taken together, these data indicate that LT exposure drives a shift in epithelial composition by promoting proliferation and suppressing differentiation.

**Table 2.**
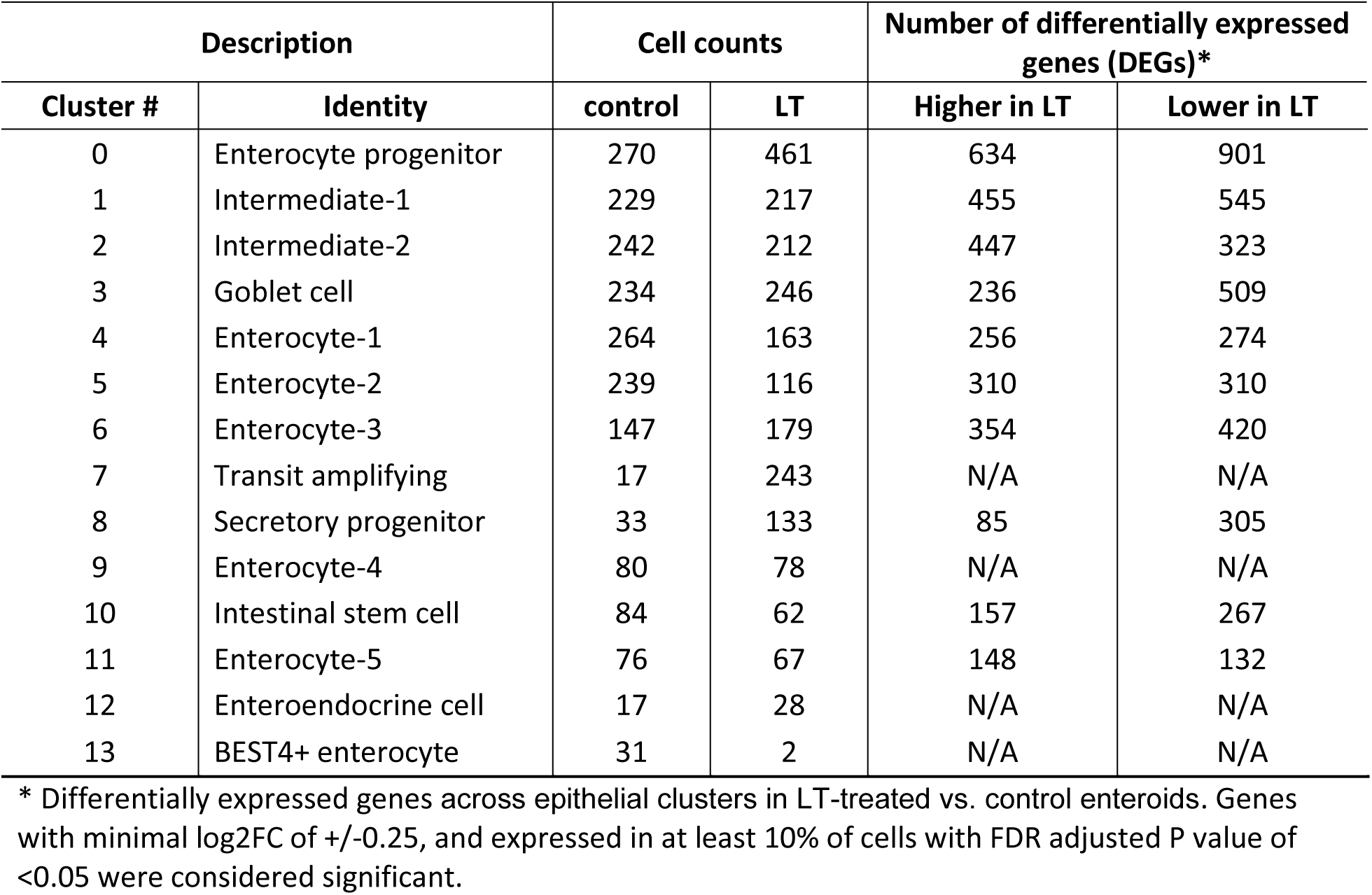
Total number of differentially expressed genes per cluster following LT treatment.

| Description |  | Cell counts |  | Number of differentially expressed genes (DEGs)* |  |
| --- | --- | --- | --- | --- | --- |
| Cluster # | Identity | control | LT | Higher in LT | Lower in LT |
| 0 | Enterocyte progenitor | 270 | 461 | 634 | 901 |
| 1 | Intermediate-1 | 229 | 217 | 455 | 545 |
| 2 | Intermediate-2 | 242 | 212 | 447 | 323 |
| 3 | Goblet cell | 234 | 246 | 236 | 509 |
| 4 | Enterocyte-1 | 264 | 163 | 256 | 274 |
| 5 | Enterocyte-2 | 239 | 116 | 310 | 310 |
| 6 | Enterocyte-3 | 147 | 179 | 354 | 420 |
| 7 | Transit amplifying | 17 | 243 | N/A | N/A |
| 8 | Secretory progenitor | 33 | 133 | 85 | 305 |
| 9 | Enterocyte-4 | 80 | 78 | N/A | N/A |
| 10 | Intestinal stem cell | 84 | 62 | 157 | 267 |
| 11 | Enterocyte-5 | 76 | 67 | 148 | 132 |
| 12 | Enteroendocrine cell | 17 | 28 | N/A | N/A |
| 13 | BEST4+ enterocyte | 31 | 2 | N/A | N/A |

\* Differentially expressed genes across epithelial clusters in LT-treated vs. control enteroids. Genes with minimal log2FC of +/-0.25, and expressed in at least 10% of cells with FDR adjusted P value of <0.05 were considered significant.

### Heat-labile toxin promotes intestinal epithelial cell proliferation and reshapes cell cycle dynamics across epithelial cell clusters

Because proliferation kinetics directly influence epithelial cell-type composition^44^, we next examined how LT affects the cell cycle status of intestinal epithelial cells. Using canonical gene sets for G1, S, and G2/M phases^46^ (Supplementary figure 5a), we assigned cell cycle stages to individual cells in LT-treated and control enteroids. UMAP projections revealed a marked shift in overall cell cycle distribution upon LT exposure (Figure 4a). Specifically, LT-treated cells showed a significant expansion of S-phase (p = 2.21 x 10^-5^, χ^2^) and G2/M-phase populations (p <1 x 10^-16^), accompanied by a reduction in G1-phase cells (Table 3).

**Figure 4:**
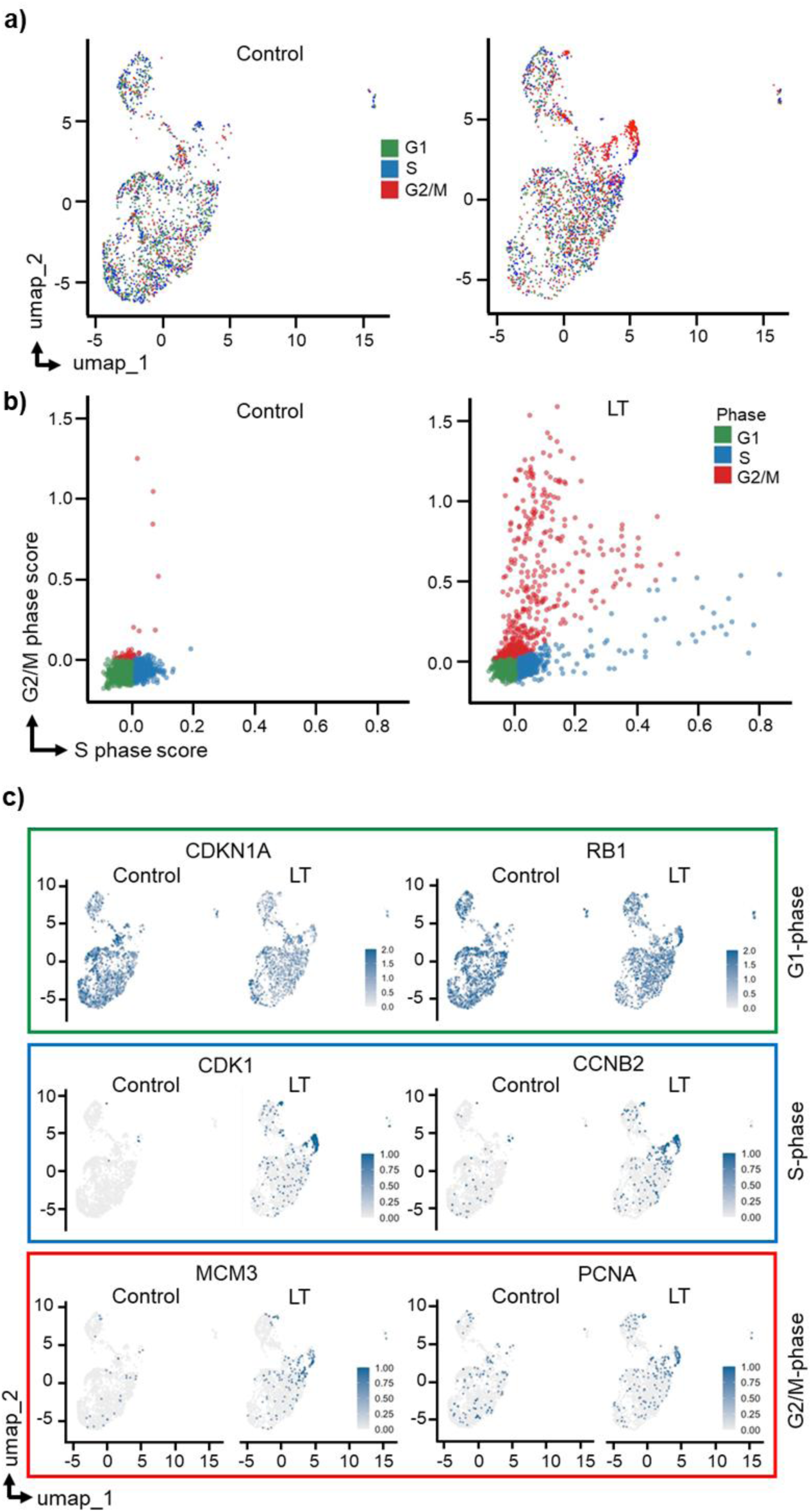
LT alters epithelial cell cycle dynamics. **a**) UMAP projections of single-cell transcriptomes from control and LT-treated human ileal enteroids, colored by assigned cell cycle phase based on canonical marker gene expression (G1, S, G2/M). **b**) XY scatterplots of individual cell S-phase and G2/M-phase scores derived from gene expression signatures. **c**) UMAP feature plots highlighting expression of representative phase-specific marker genes across conditions. G1-associated genes (RB1, CDKN1A), S-phase associated genes (PCNA, MCM3) and G2/M-phase associated genes (CCNB2, CDK1). Color scales represent normalized expression.

**Table 3.** Cell cycle phase distribution in control and LT-treated enteroids.

| Clusters | Phase | Cell count (%) |  | Chi<sup>2</sup>_statistic | P value |
| --- | --- | --- | --- | --- | --- |
|  |  | Control | LT |  |  |
| all combined | G1 | 1303 (66.4) | 1031 (46.8) | 162.2 | < 1e-16 |
|  | S | 602 (33.7) | 546 (24.7) | 18 | 2.21E-05 |
|  | G2/M | 58 (2.9) | 630 (28.5) | 492.01 | < 1e-16 |
| #0 | G1 | 197 (73.0) | 221 (47.9) | 61.06 | <0.00001 |
|  | G2M | 10 (3.7) | 107 (23.2) |  |  |
|  | S | 63 (23.3) | 133 (28.9) |  |  |
| #1 | G1 | 144 (62.9) | 104 (47.9) | 43.74 | <0.00001 |
|  | G2M | 6 (2.6) | 51 (23.5) |  |  |
|  | S | 79 (34.5) | 62 (28.6) |  |  |
| #2 | G1 | 137 (56.6) | 139 (65.6) | 37.596 | <0.00001 |
|  | G2M | 7 (2.9) | 32 (15.1) |  |  |
|  | S | 98 (40.5) | 41 (19.3) |  |  |
| #3 | G1 | 187 (79.9) | 147 (59.8) | 53.1702 | <0.00001 |
|  | G2M | 3 (1.3) | 57 (23.2) |  |  |
|  | S | 44 (18.8) | 42 (17.1) |  |  |
| #4 | G1 | 176 (66.7) | 82 (50.3) | 23.0313 | <0.00001 |
|  | G2M | 1 (0.4) | 12 (7.4) |  |  |
|  | S | 87 (33.0) | 69 (42.3) |  |  |
| #5 | G1 | 165 (69.0) | 67 (57.8) | 21.4174 | 0.000022 |
|  | G2M | 2 (0.8) | 13 (11.2) |  |  |
|  | S | 72 (30.1) | 36 (31.0) |  |  |
| #6 | G1 | 95 (64.6) | 110 (61.5) | 10.847 | 0.004412 |
|  | G2M | 7 (4.8) | 28 (15.6) |  |  |
|  | S | 45 (30.6) | 41 (22.9) |  |  |
| #7 | G1 | 7 (41.2) | 1 (0.4) | 90.4198 | <0.00001 |
|  | G2M | 7 (41.2) | 214 (88.1) |  |  |
|  | S | 3 (17.6) | 28 (11.5) |  |  |
| #8 | G1 | 18 (54.5) | 37 (27.8) | 11.3691 | 0.0034 |
|  | G2M | 5 (15.2) | 58 (43.6) |  |  |
|  | S | 10 (30.3) | 38 (28.6) |  |  |
| #9 | G1 | 48 (60.0) | 41 (52.6) | 5.5985 | 0.061 |
|  | G2M | 4 (5.0) | 13 (16.7) |  |  |
|  | S | 28 (35.0) | 24 (30.8) |  |  |
| #10 | G1 | 68 (81.0) | 22 (35.5) | 38.4988 | <0.00001 |
|  | G2M | 3 (3.6) | 25 (40.3) |  |  |
|  | S | 13 (15.5) | 15 (24.2) |  |  |
| #11 | G1 | 44 (57.9) | 39 (58.2) | 19.4254 | 0.000061 |
|  | G2M | 1 (1.3) | 15 (22.4) |  |  |
|  | S | 31 (40.8) | 13 (19.4) |  |  |
| #12 | G1 | 15 (88.2) | 20 (71.4) | 1.4768 | 0.47787 |
|  | S | NF* | 5 (17.9) |  |  |
|  | G1 | 2 (11.8) | 3 (10.7) |  |  |
| #13 | G1 | 2 (6.5) | 1 (50.0) | 5.4403 | 0.065863 |
|  | G2M | 2 (6.5) | NF* |  |  |
|  | S | 27 (87.1) | 1 (50.0) |  |  |

\*NF-not found

Cell cycle phase scores further confirmed these shifts, with elevated expression of S-phase and G2/M gene sets in LT-treated enteroids (Figure 4b). Consistent with this, we observed differential expression of key phase-specific cell cycle regulators, including RB1 and CDKN1A (G1-phase), PCNA and MCM3 (S-phase), and CCNB2 and CDK1 (G2/M-phase) (Figure 4c). LT exposure also upregulated stem and progenitor markers such as CD24, CD44, and OLFM4, reinforcing the expansion of the undifferentiated, proliferative compartment (Supplementary figure 5b).

Compartmentalization of intestinal cell proliferation is critical for epithelial maintenance and function with higher proliferative activity at the crypt zone and no proliferation at the tips of villi. Analysis of cell cycle dynamics across transcriptionally defined clusters revealed a widespread reduction in G1-phase cells and robust enrichment of G2/M-phase cells across epithelial subpopulations (Figure 5a; Table 3), indicating that LT alters cell cycle dynamics across the entire epithelial landscape.

**Figure 5:**
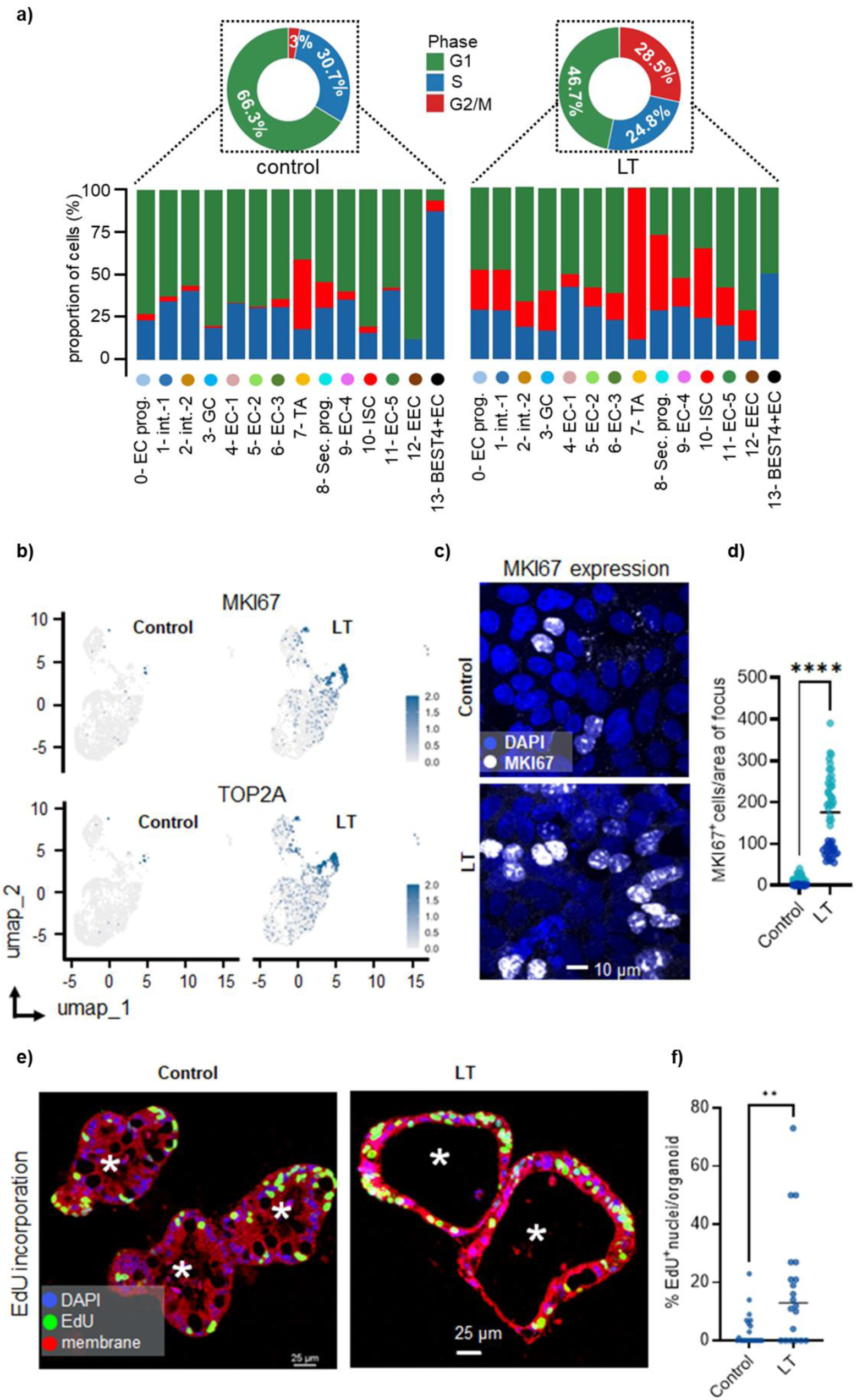
LT affects cell cycle progression across intestinal epithelial lineages and promotes epithelial proliferation. **a**) Donut charts represent the global distribution of cells across cell cycle phases in control and LT-treated enteroids. Bar plots below show the proportion of cells in G1 (green), S (blue), and G2/M (red) phases across transcriptionally defined cell clusters (0–13) in control and LT-treated enteroids. **b**) UMAP feature plots showing expression of proliferation markers MKI67 (top) and TOP2A (bottom) in control (left) and LT-treated (right) human ileal enteroids. The color scale indicates normalized expression levels. **c**) Representative confocal images of MKI67 (white) and nuclei (DAPI, blue) in control (left) and LT-treated (right) enteroids. Scale bar, 10 μm. **d**) Quantification of MKI67⁺ cells per field of view. Each point represents the number of MKI67⁺ nuclei in an individual field; the horizontal line denotes the mean. Data are from two independent experiments (≥50 fields of view total). Statistical significance was determined by a two-tailed, nonparametric Mann–Whitney test (****p < 0.0001). **e**) Representative confocal images of control (left) and LT-treated (right) enteroids labeled with EdU (green) for DNA synthesis, DAPI (blue) for nuclei, and a membrane stain (red). White asterisks mark the luminal space (equivalent to intestinal lumen) in enteroids. Off note, LT-treated enteroids showed a marked increase in luminal area, consistent with LT-induced fluid secretion. Scale bar, 25 μm. **f**) Quantification of EdU⁺ nuclei per enteroid (n = 25 enteroids per treatment). Each point represents the number of EdU^+^ nuclei per individual enteroid; the horizontal line denotes the mean. Statistical significance was determined by a two-tailed, nonparametric Mann–Whitney test (**p < 0.01).

To further support the observed shift in cell cycle dynamics, we analyzed the expression of canonical proliferation markers. LT-treated enteroids exhibited strong upregulation of MKI67 and TOP2A, two well-established indicators of active proliferation^47^ (Figure 5b). To validate these transcriptomic findings at the protein level, we performed phenotypic assays using polarized epithelial monolayers. Enteroids were cultured on Transwell filters to establish apical–basolateral polarity and treated with LT during the differentiation phase. Confocal immunofluorescence microscopy revealed a significant increase in MKI67⁺ nuclei in LT-treated monolayers compared to controls (Figure 5c-d), confirming enhanced proliferative activity at the protein level.

To functionally validate these observations, we utilized 5-ethynyl-2’-deoxyuridine (EdU) incorporation assay in 3D culture of enteroids treated with LT. Following differentiation, both control and LT-treated enteroids were incubated with EdU. LT treatment led to significantly higher incorporation of EdU (p<0.01), indicating enhanced DNA synthesis and entry into S phase (Figure 5e–f). Collectively, these findings indicate that LT skews epithelial homeostasis by promoting a proliferative, progenitor-dominated state across the intestinal epithelium.

### Heat-labile toxin suppresses epithelial differentiation and impairs functional maturation

Given this marked shift away from differentiated lineages, we next investigated whether LT-induced disruption of differentiation affects epithelial maturation. Villin, a structural component of apical microvilli, and intestinal alkaline phosphatase (ALPI), a brush-border enzyme, serve as canonical markers of mature enterocytes^12,48^. To assess the functional consequences of LT-induced differentiation defects, we examined villin and ALPI protein expression in membrane and cytosolic fractions, respectively, from control and LT-treated enteroids. Consistent with the transcriptional changes (Supplementary Figure 6a), immunoblot analysis showed significantly reduced expression of villin, and alkaline phosphatase (ALPI), in LT-treated enteroids compared to controls (Figure 6a–b), suggesting impaired epithelial maturation at the protein level.

**Figure 6.**
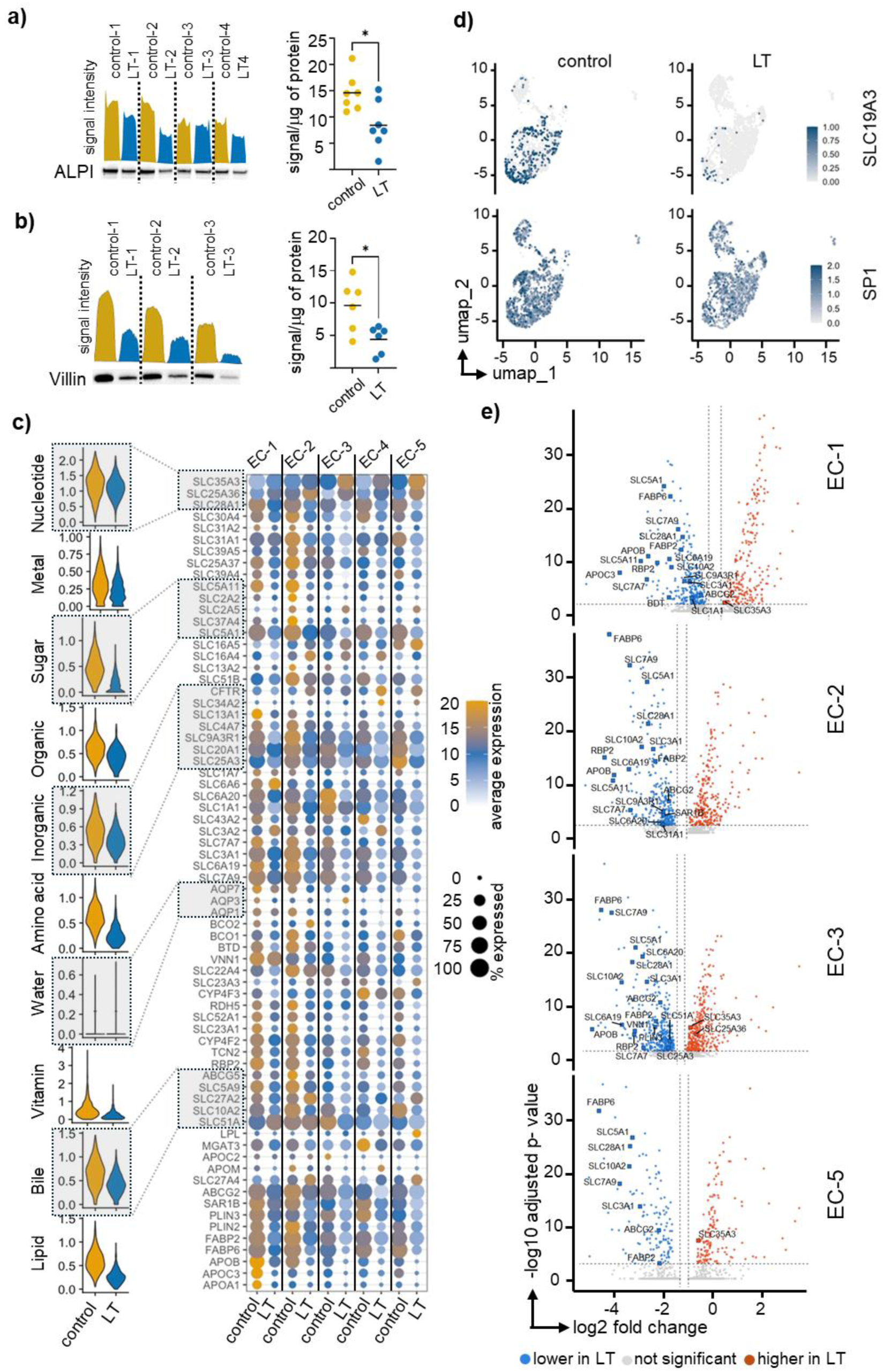
LT affects enterocyte maturation and disrupts nutrient transporter expression. **a**) Western blot analysis of ALPI (intestinal alkaline phosphatase enzyme) expression in control and LT-treated enteroids. Representative blot and corresponding densitometric histograms generated using ImageJ (left). Quantification of normalized ALPI expression (right). Each data point represents an individual replicate (n = 7 per group). Horizontal lines on the plot indicate mean. Statistical significance was determined using a non-parametric Mann-Whitney test (*p < 0.05, two-tailed). **b**) Western blot analysis of Villin (structural protein) expression in control and LT-treated enteroids. Representative blot and corresponding densitometric histograms generated using ImageJ (left). Quantification of normalized Villin expression (right). Each data point represents an individual replicate (n = 6 per group). Horizontal lines on the plot indicate mean. Statistical significance was determined using a non-parametric Mann-Whitney test (*p < 0.05, two-tailed). **c**) Analysis of nutrient transporter genes across enterocyte subclusters (EC-1 to EC-5). Data were derived from scRNA-seq of control and LT-treated enteroids. Violin plots showing distributions of mean expression of major nutrient transporter gene families (nucleotide, metal, sugar, organic, inorganic, amino acid, water, vitamin, bile, lipid). Dot plot illustrating expression patterns of specific genes involved in nutrient absorption and transport. The color scale indicates the average expression, and the dot size indicates proportion of cells expressing individual transporter genes across five enterocyte subclusters (EC-1 to EC-5). **d**) UMAP feature plots showing SLC19A3 thiamine transporter and SP1 regulator gene expression across cell clusters. **e**) Volcano plots showing differential expression of nutrient transporter genes within enterocyte subclusters EC-1 through EC-5. The x-axis indicates log₂ fold-change (LT vs. control), and the y-axis indicates –log₁₀ adjusted p-value. Genes significantly upregulated in LT-treated enteroids are shown in red, significantly downregulated genes in blue, and non-significant changes in gray. Selected transporters with significant changes are labeled. Statistical testing was performed using a Wilcoxon rank-sum test with Benjamini–Hochberg correction for multiple comparisons; adjusted p < 0.05 was considered significant. EC, enterocyte.

To further investigate the impact of LT on the functional maturation of the epithelia, we examined the expression of key solute carrier (SLC) genes responsible for nutrient transport. Although we observed robust expression of multiple SLC family members in control enteroids, LT-treated enteroids exhibited marked repression of these genes across the epithelial cell clusters (Supplementary figure 6b). Notably, LT diminished the expression of SLC28A1 (nucleoside), SLC23A1 (vitamin C), SLC30A1/30A4 (zinc), SLC7A7 (amino acids), SLC5A11 (myo-inositol), SLC19A3 (thiamine), SLC11A2 (iron), SLC17A4 (glutamate), SLC15A1 (di/tripeptides), and other transporters (Supplementary figure 7), consistent with a depletion of functionally mature enterocytes.

Because nutrient absorption is a key functional hallmark of mature enterocytes, we next analyzed the transcriptional consequences of LT treatment on nutrient transporter expression across five transcriptionally defined enterocyte subclusters (EC1–EC5). LT exposure led to a global reduction in transporter gene expression scores across multiple categories, including sugar, amino acid, water, bile acid, lipid, and vitamin transport (Figure 6c, left). This was further supported by reduced average expressions and cell-level prevalence of multiple solute carrier (SLC) family genes^12^ in LT-treated cells compared to controls (Figure 6c, right).

Consistent with these observations, prior bulk RNA-seq studies have shown that LT suppresses SLC19A3, a thiamine transporter, along with its upstream regulator SP1, leading to impaired thiamine uptake^49^. Our single-cell RNA-seq data recapitulated this finding, with decreased expression of both SLC19A3 and SP1 in LT-treated enteroids (Figure 6d). Additionally, differential gene expression analysis of enterocyte clusters revealed significant downregulation of other key absorptive genes, including SLC5A1 (sodium/glucose cotransporter 1), SLC10A2 (ileal sodium/bile acid cotransporter), SLC28A1 (sodium/nucleoside cotransporter 1), FABP6 (fatty acid transporter), and APOB (fatty acid transporter), confirming suppression of essential nutrient acquisition pathways (Figure 6e).

Altogether, these studies suggest that LT exerts a profound effect on the developing intestinal epithelia. The LT-driven proliferative response coincided with transcriptional activation of WNT/β-catenin target genes and nuclear accumulation of β-catenin, consistent with canonical WNT pathway activation. By impacting major transcriptional pathways, including those orchestrated by β-catenin, LT disrupts epithelial differentiation and subsequent maturation.

## Discussion

ETEC infections have repeatedly been linked to long-term sequelae including malnutrition and impaired growth, significantly compounding the morbidity burden beyond the acute infection ^5,8,15,16,50,51^. While LT-producing ETEC have specifically been linked to malnutrition among young children^50^, the precise molecular mechanisms underlying links between ETEC and nutritional sequelae remain incompletely understood. The present studies suggest that LT may alter the trajectory of epithelial cell maturation, stalling the development of mature absorptive enterocytes. This disruption of normal epithelial homeostasis parallels earlier observations that LT also interferes with transcriptional programs that govern the biogenesis of enterocyte microvilli, comprising the major surface for nutrient absorption, along with LT modulation of a multitude of nutrient transporters^52^. The collective impacts of LT on the architecture and function of absorptive intestinal epithelia likely contribute significantly to the nutritional deficits that accompany ETEC infections in young children.

Here we show that the heat-labile enterotoxin reprograms intestinal epithelia by subversion of the fundamental regulation of stem cell maintenance, proliferation, and differentiation^18,53,54^ governed by β-catenin. While LT is classically known for inducing diarrhea via cAMP-mediated chloride secretion^55,56^, its broader impact on intestinal development has remained poorly characterized. Here, we show that LT activates β-catenin signaling independent of WNT by stabilizing β-catenin through PKA-mediated phosphorylation of β-catenin and components of its degradation complex^32,57,58^. The resulting enhanced and aberrant β-catenin signaling ultimately reprograms homeostatic maturation of intestinal epithelia.

The intricate nature of cAMP and PKA signaling is reflected in complex array of changes that follow exposure to heat-heat labile toxin. Mechanistically, LT modulates transcriptional programs by PKA-mediated phosphorylation of a family of cAMP-responsive nuclear factors including the cAMP response element binding protein (CREB) and the cAMP response element modulator (CREM), permitting their interaction with transcriptional co-activators including CREB-binding protein (CBP) as they bind cooperatively to short DNA consensus sequences known as cAMP response elements (CRE)^59^ in the promoter regions of target genes. Intriguingly, genetic studies in Bangladeshi children identified CREM polymorphisms associated with increased susceptibility to diarrhea and reduced weight-for-age^60^, potentially supporting the involvement of cAMP signaling in nutritional sequelae.

The cAMP pathway is also at the center of complex regulatory cascades involving multiple transcription factors in addition to CREB and CREM that are also modulated by PKA^59^. These include the transcriptional regulators HNF4A and SMAD4, both essential for enterocyte differentiation and known to be sensitive to PKA-dependent phosphorylation^61,62^ ^63-65^. Beyond direct transcriptional effects, cAMP-driven activation of co-regulators such as CBP and PHF2 can induce histone acetylation and methylation, also raising the possibility that LT reinforces impaired epithelial maturation through epigenetic remodeling^62^.

Our data supports the view that LT activates β-catenin signaling to impair terminal enterocyte differentiation. Under homeostatic conditions, β-catenin regulates the balance between stem cell renewal and lineage commitment in the crypt-villus axis^21,86-88^. Disruption of this axis, such as through loss of WNT antagonists like APC, leads to crypt expansion and impaired epithelial maturation ^21,66,67^. Our findings align with this paradigm, showing that LT mimics hyperactive WNT signaling to expand progenitor and transit-amplifying populations while depleting differentiated lineages, including mature enterocytes and BEST4⁺ absorptive cells^68^. This imbalance is characteristic of a poorly differentiated epithelium and closely mirrors the pathological features of environmental enteric dysfunction (EED), characterized by villus blunting, crypt expansion, and malabsorption^69,70^.

The depletion of mature absorptive cells and downregulation of nutrient transporter genes in LT-treated enteroids recapitulate key features of immature epithelia. Epithelial maturation involves functional enrichment of nutrient absorption capacity in villus tip cells, where genes required for brush border formation and fat absorption are upregulated in differentiated villus enterocytes^12^. However, LT-treated epithelia exhibit reduced expression of ALPI, and VIL1, canonical markers of enterocyte maturation, as well as transcriptional repression of multiple solute carrier family genes essential for absorptive functions of nutrient, bile acid, and vitamin transport. The failure to achieve mature enterocyte identity is particularly relevant in the context of EED, where altered bile acid metabolism and nutrient malabsorption are linked to stunted growth. Notably, a study in undernourished Pakistani infants found that disrupted bile acid homeostasis correlated with EED histopathology and growth faltering^71^. LT-induced repression of bile acid transporters (SLC10A2, FABP6) may offer a plausible mechanistic link to these clinical observations.

This concept is further supported by recent animal studies, where bile acid accumulation and NAD⁺ deficiency induced by a low-protein diet led to villus atrophy and epithelial loss, phenotypes that were reversed by dual supplementation with NAD⁺ precursors and bile acid sequestrants ^72^. Our findings raise the possibility that LT could act synergistically with nutritional stressors to maintain an immature epithelial state, exacerbating malabsorption and stunting in vulnerable children. The ability of LT to disrupt intestinal maturation and impair nutrient uptake underscores the pathogenic potential of ETEC beyond acute diarrhea.

From a broader perspective, this work positions LT as a potential molecular link between ETEC infection and long-term intestinal dysfunction. The sustained shift toward a proliferative, undifferentiated epithelial state, driven by toxin-mediated signaling, mirrors intestinal phenotypes observed in chronically infected or malnourished children, where impaired differentiation contributes to long-term functional deficits^8,16,17,73^. These findings are especially concerning in the context of repeated or chronic ETEC exposure, which is common in low-resource settings and has been linked to growth faltering in children even in the absence of overt diarrhea^15,50,51,74-78^.

While this study provides mechanistic insight into LT-induced epithelial reprogramming, several limitations should be noted. First, the findings are based on *in vitro* enteroid models, which lack stromal, immune, and microbial components that shape epithelial responses *in vivo.* Second, although we identify β-catenin signaling as a key mediator, the specific upstream effectors connecting LT-induced cAMP production to transcriptional reprogramming remain to be fully defined. In addition to LT, many ETEC make heat-stable toxins (ST) which engage guanylate cyclase C (GC-C) to stimulate production of cGMP and activation of protein kinase G (PKG). Although ST has not been thoroughly evaluated in the context of EED, ST and other GC-C agonists are known to impact cell cycle progression and modulate epithelial proliferation and differentiation^79-81^.

Understanding how these common enterotoxins contribute to pathogenesis of intestinal dysfunction that follow ETEC infections can inform the rational design of vaccines that protect against acute illness as well as the tremendous burden of sequelae^52,82^. This work advances our understanding of how ETEC contributes to long-term intestinal dysfunction and malnutrition and offers a framework for developing strategies to mitigate the persistent sequelae of enteric infections in vulnerable populations.

## Materials and methods

All experiments were conducted in accordance with relevant ethical guidelines, and study protocols were approved by the Institutional Biosafety Committee (IBC), and Institutional Review Board (IRB) at Washington University School of Medicine in St. Louis.

### Growth and maintenance of small intestinal enteroid cell lines

Hu235D derived from human small intestinal (ileal) stem cells were obtained from the Precision Animal Models and Organoids Core ([PAMOC](https://ddrcc.wustl.edu/scientific-cores/pamoc/)) maintained by the Washington University Digestive Diseases Research Core Center (DDRCC), and were propagated as previously described^83^. Briefly, crypt intestinal stem cells maintained in liquid nitrogen were thawed, resuspended in Cultrex (Cultrex BME Type II, Catalog# 3532-010-02, Biotechne) and transferred to wells in a 48-well plate. Cells were grown for 3-4 days in growth media containing 50% L-WRN conditioned media derived from the L-WRN cell line (provided by the POMAC), a 1:1 mixture of primary media [Advanced DMEM/F12, Invitrogen 12634028, supplemented with fetal bovine serum (FBS, 20%), 2 mM L-glutamine, 100 U/ml penicillin, 0.1 mg/ml streptomycin], 10 µM ROCK inhibitor Y-27632 (Tocris Bioscience) and 10 μM TGFBR1 inhibitor SB 431542 (Tocris Bioscience)^83^,^84^.

For RNA-seq analysis, cells were grown in Cultex for 4 days for enteroid formation, diluted 1:2 in Cultrex to reduce enteroid density, replated and grew for 3 more days in growth media. To allow differentiation, enteroids were then cultured for additional 3 days in differentiation media (5% L-WRN conditioned media supplemented with 10 µM ROCK inhibitor). Enteroids were treated with heat-labile toxin (100ng/ml) either during the differentiation period (LT), or PBS treated (control). For polarized enteroid monolayers, following trypsinization cells were seeded onto Transwell filters (Corning) coated with type IV human collagen (Sigma) and grew in growth media for confluent monolayer. After confluency growth, media was replaced with differentiation media to promote differentiation.

### Quantitative RT-PCR

Quantitative real-time PCR was conducted on a QuantStudio 3 detection system (Applied Biosystems) using Fast SYBR Green master mix (Applied Biosystems/Thermo Fisher). Specificity of amplification for each sample was confirmed by dissociation curve analysis, and PCR product size was verified through agarose gel electrophoresis. For transcript amplification from small intestinal enteroids, previously described methods were followed, with relative gene expression normalized to GAPDH. All primers used are provided in Supplementary table 1.

### Fluorescence microscopy

For fluorescence microscopy, polarized enteroid monolayers were fixed in 2% paraformaldehyde for 30 minutes at 37°C, followed by an additional 30 minutes at room temperature. After fixation, the monolayers were washed three times with PBS and blocked with 2% BSA-PBS to minimize non-specific binding. The primary antibodies (as detailed in Supplementary table 2) used for labeling specific proteins were diluted in 2% BSA-PBS and incubated with the monolayers for 1 hour at room temperature. Fluorescent dye-conjugated secondary antibodies were then diluted in 2% BSA-PBS and applied to the monolayers for 45 minutes at room temperature. Cell membranes and nuclei were stained using CellMask (1:2000 dilution, Invitrogen) and DAPI (1:1000 dilution, Invitrogen), respectively. Slides were washed 3x with PBS and coverslipped with Prolong Gold antifade reagent (Invitrogen). Images were captured and analyzed using a Nikon C2 confocal microscope equipped with NIS-Elements AR 5.11.01 software (Nikon).

### Construction of TCF/LEF reporter enteroid lines

HEK293T cells (ATCC CRL-3216) were propagated at 37°C with 5% CO₂ in Dulbecco’s Modified Eagle’s Medium (DMEM) supplemented with 10% heat-inactivated fetal bovine serum (FBS) and 2 mM L-glutamine in T25 flasks. For 7TFP containing lentivirus production, cells were co-transfected with 5 µg of p7TFP, 5 µg of psPAX2, and 3 µg of pMD2.G plasmids (see Supplementary table 3) in a 500 µl of culture medium with 39 µl of FuGENE HD transfection reagent (Promega). Media was changed following 18 h post transfection. Two days post transfection, the 7TFP-lentivirus–containing supernatant was collected, clarified by centrifugation at 3,000 rpm for 15 minutes, and concentrated by adding polyethylene glycol (PEG) 8000 at a 1:3 ratio (PEG:supernatant). The mixture was rotated overnight at 4°C and centrifuged at 1,600 × g for 1 hour. The resulting viral pellet was resuspended in 50% conditioned medium and used to transduce Hu235D human small intestinal enteroids, in the presence of polybrene (final concentration: 8 µg/ml). After 8 hours at 37°C, cells were pelleted, resuspended in fresh Cultrex, and cultured for 3 days. Transduced cells were selected using puromycin (2.5 µg/ml) for p7TFP-expressing enteroids (Hu235D::7TFP) and sorted in liquid nitrogen.

### 7TCF/LEF Luciferase assays

The Hu235D::7TFP cell line was resuspended in 50% L-WRN conditioned media enumerated (Countess FL, ThermoFisher Scientific), and 35,000 live cells added in a total volume of 100 µl per well of a 96 well plate (TPP, flat bottom, Cat# 92096) pre-coated with 3% Cultrex. After 48 hours growth at 5% CO_2_, 37°C, media was replaced with 5% L-WRN + Y-27632 to encourage differentiation. After 2 days, cells in respective wells (∼95% confluency) were treated with toxins as indicated in primary media only. After equilibration to room temperature, 5’-fluoroluciferin reagent (ONE-Glo EX, E8110 Promega) was added to wells and plates shaken at 200 rpm for 10 minutes. 120 µl from each well was then transferred to corresponding wells of an opaque white 96 well plate (Microlite 1+, ThermoFisher 7417) and luminescence measured (Synergy H1, BioTek). All treatments were performed at least in triplicate, repeated at least three times.

### Subcellular fractionation and immunoblotting

To isolate nuclear and cytoplasmic fractions, subcellular fractionation of human ileal enteroids was performed using the NE-PER™ Nuclear and Cytoplasmic Extraction Kit (Thermo Fisher Scientific, #78833), following the manufacturer’s protocol with minor adaptations. Enteroids (∼1 × 10⁶ cells per condition) were collected after treatment (e.g., with heat-labile toxin or PBS control), washed twice in ice-cold PBS, and pelleted by centrifugation at 500 × g for 5 min at 4°C. Cytoplasmic and nuclear extracts were sequentially isolated using the provided CER I, CER II, and NER buffers supplemented with Halt™ protease and phosphatase inhibitor cocktail (Thermo Fisher Scientific, #78440). After incubation with CER I and CER II to lyse the cytoplasmic membrane, lysates were centrifuged at 16,000 × g for 5 min to separate the supernatant for cytoplasmic fraction. The pellet was then resuspended in NER buffer and incubated with vigorous vortexing every 10 min over a 40-min period. The nuclear fraction was collected by centrifugation at 16,000 × g for 10 min. Protein concentrations were measured using a BCA assay (Thermo Fisher Scientific), and fraction purity was confirmed by immunoblotting for cytoplasmic (GAPDH) and nuclear (Lamin b1) markers. For immunoblotting, 10–30 µg of protein per sample was mixed with Laemmli sample buffer containing 100 mM DTT, boiled at 95°C for 10 min, and resolved on 4–12% SDS-PAGE gels. Proteins were transferred to nitrocellulose membranes (100 V, 1 h) and blocked in 5% non-fat milk in PBS-Tween (0.1%). Membranes were incubated overnight at 4°C with primary antibodies at 1:1000 dilutions (Supplementary table 2). After washing three times with PBST, HRP-conjugated secondary antibodies (1:5,000) were applied for 1 h at room temperature. Signals were developed using Clarity ECL substrate (BioRAD) and visualized with a c600 imager (azure Biosystems). Band intensities were quantified using ImageJ and normalized to total protein across biological replicates.

### Single nuclei isolation for RNA sequencing

Following treatment, enteroids were collected in ice-cold PBS supplemented with 0.5 mM EDTA, resuspended, and centrifuged at 200g for 5 minutes. For single-cell preparation, the cell pellets were resuspended in TrypLE (Invitrogen) and incubated at 37°C for 8 minutes to obtain a single-cell suspension. Cell debris and large clumps were removed by passing the single-cell suspension through a 40 μm filter, followed by centrifugation at 250g for 5 minutes. Prior to nuclei isolation, the cells were washed once with ice-cold PBS supplemented with 0.04% BSA and counted to obtain 1 million cells. Single nuclei were isolated following the protocol for nuclei isolation from fresh cells (Protocol CG000365 Rev C, 10X Genomics). Briefly, 100 μl of chilled Lysis Buffer was added to the cell pellet, mixed by pipetting 10 times, incubated for 5 minutes on ice, and washed with 1 ml of Wash Buffer. Nuclei were pelleted by centrifugation at 500g for 5 minutes at 4°C, washed two more times, then resuspended in Nuclei Buffer and immediately processed for sequencing.

### cDNA library preparation and sequencing

For single-cell RNA sequencing together with single-cell ATAC sequencing, Multiome 3v3.1 GEX and ATAC libraries were prepared by using the Chromium Next GEM Single Cell Multiome ATAC + Gene Expression Reagent Kits following the recommended protocol (User Guide, CG000338, 10X Genomics). Samples were prepared on the 10x Genomics platform using the Chromium Next GEM Single Cell Multiome ATAC + Gene Expression Reagent Bundle (16 rxns, PN-1000283), Chromium Next GEM Chip J Single Cell Kit (48 rxns, PN-1000234), Single Index Kit N Set A (96 rxns, PN-1000212, for ATAC), and Dual Index Kit TT Set A (96 rxns, PN-1000215, for 3v3.1 GEX). The concentration of each library was determined by qPCR using the KAPA Library Quantification Kit (qPCR MasterMix cat#7959478001, and Library Quantification Kit, Cat# 7960085001, Roche Diagnostics System), following the manufacturer’s protocol, to achieve appropriate cluster counts for the Illumina NovaSeq6000 instrument. GEX libraries were pooled and run on 0.0192 of a NovaSeq X Plus 25B flow cell using a 28x10x10x150 sequencing recipe, as per the manufacturer’s protocol, with a target coverage of 500M read-pairs per sample. ATAC libraries were pooled and run on 0.0115 of a NovaSeq X Plus 25B flow cell with a 51x8x16x51 sequencing recipe, following the manufacturer’s protocol, aiming for a target coverage of 300M read-pairs per sample.

### Data processing

Single-cell RNA-seq and ATAC-seq data were processed using the 10x Genomics Cell Ranger ARC pipeline (version 2.0.2). For alignment and quantification of gene expression data, raw sequencing reads were processed using the default parameters provided by the 10x Genomics software suite. The standard reference genome and annotations used were based on the refdata-cellranger-arc-GRCh38-2020-A-2.0.0 build provided by 10x Genomics. Briefly, the Illumina sequencer’s base call files (BCLs) for each flow cell directory (ATAC or Gene Expression) were demultiplexed using the Illumina bcl2fastq software to generate FASTQ files. These reads were then aligned to the GRCh38 human reference genome, and unique molecular identifier (UMI) counts were obtained using the 10x Cell Ranger toolkit. Barcode selection and filtering for single-cell droplets were performed according to the Cell Ranger pipeline^11,27^

### Data filtering and normalization

Single-cell Multiome libraries were used for initial quality control and cell filtering. For the present study, downstream analyses were performed exclusively on the RNA-seq data, which is the focus of this manuscript. Analyses of the ATAC-seq data will be reported separately. UMI count matrices for single cells were analyzed in Seurat (v5.0)^45^. To ensure data quality, cells with fewer than 1,000 RNA counts or more than 25,000 RNA counts were removed, along with cells where more than 50% of reads were mapped to mitochondrial genes, thereby excluding potential low-quality cells, debris, and likely multiplets. To further refine the dataset, doublets were identified and removed using DoubletFinder^85^ based on the expected doublet rate and model homotypic interactions, ensuring more accurate cell classification. Following these steps, we applied the SCTransform method, a regularized negative binomial regression approach that corrects for sequencing depth and other technical variations while preserving biological heterogeneity in the dataset^86^. The output from SCTransform was used to identify highly variable features, which were then used for downstream analysis.

### Marker gene identification for cell clustering

To identify marker genes across all clusters, we used the FindAllMarkers function in Seurat. Genes were designated as marker genes for a given cluster if they were significantly upregulated in that cluster (Wilcoxon rank-sum test p < 0.05), expressed in at least 25% of cells within the cluster, and showed a minimum log fold change of 0.25 (indicating approximately 19% higher expression) relative to other clusters. Expression heatmaps of the top 10 signature genes for each cluster are shown in Figure 3b, and all marker genes names and statistics for their enrichment are provided for every cluster in **Supplementary dataset 1.**

### Differential gene expression analysis

To identify gene expression variability and cell-specific responses to treatment, differential gene expression within individual cell clusters was analyzed. Differentially expressed genes (DEGs) were identified using Seurat’s FindMarkers function to evaluate transcriptional changes across cell clusters following LT treatment. For each cluster, DEGs were determined using a log2-fold-change threshold of ±0.25 to capture biologically relevant differences, and considering only genes expressed in at least 30% of cells within each group. The Wilcoxon rank-sum test was used for statistical comparisons, allowing identification of both up- and down-regulated genes and providing a comprehensive view of treatment-induced transcriptional changes. The list of all DEGs is provided in **Supplementary dataset 2**.

### Cell cycle analysis

Each cell was annotated for its cell cycle stage using the CellCycleScoring function and the updated G1/S and G2/M cell cycle genes from the ‘cc.genes.updated.2019’ dataset in Seurat^46,87^, which evaluates the expression of stage-specific marker genes. Specifically, this function assigns each cell a quantitative score based on the expression of 54 genes related to the S phase and 43 genes associated with the G2/M phase^27106^. By comparing these scores, cells were categorized as being in G1, S, or G2/M phases. Cells with low scores in both S and G2/M markers are inferred to be in the G1 phase, indicating a quiescent or resting state, while cells with high S or G2/M scores are identified as actively cycling. Composition of cell cycle phase across different clusters is provided in supplementary source data for tables (see Supplementary table 4). For visualization, the distribution of cell cycle phases per sample and cluster was plotted using ggplot2.

### Labeling of proliferating cells

Actively proliferating cells were detected using the Click-iT EdU imaging kit (Molecular Probes, Life Technologies) according to the manufacturer’s protocol. Briefly, following toxin treatments, organoids were incubated for three hours at 37°C in tissue culture media supplemented with 10 µM EdU. Following incubation, organoids were fixed with 4% paraformaldehyde as previously described and permeabilized with 0.5% Triton X-100 for 20 minutes at room temperature. The EdU-labeled cells were detected with the Click-iT® Plus reaction cocktail, followed by counterstaining with DAPI. For EdU analysis, twenty random fields were imaged at 20× magnification, and the numbers of EdU-positive and DAPI-positive nuclei were counted. The percentage of EdU-positive nuclei was calculated as a proportion of the total nuclei count per enteroid. Each experiment was conducted in duplicate.

### Functional enrichment analysis

Ontology-independent algorithms in CompBio 2.9 (COmprehensive Multi-omics Platform for Biological InterpretatiOn) (https://gtac.wustl.edu/bioinformatics/) were used to analyze existing transcriptome data in RNAseq datasets. Enrichment analysis was carried out using WebGestalt (https://www.webgestalt.org/) against reference gene sets curated from Reactome pathways and network databases. Over-representation analysis (ORA) was performed to identify biological pathways and functional categories significantly enriched among differentially expressed genes. Genes with an adjusted *p*-value < 0.05 were selected as input. Statistical significance was determined using a hypergeometric test, with p-values adjusted for multiple testing by the Benjamini–Hochberg method (FDR < 0.05). Redundancy among enriched categories was reduced using the weighted set cover method, and results were visualized to highlight key biological processes associated with the gene sets.

### Statistical analysis

All statistical analyses were conducted using GraphPad Prism (version 10.5.0) and RStudio (version 2023.12.1), unless otherwise specified. For comparisons between two groups, the non-parametric Mann–Whitney U test was used. For comparisons among more than two groups, the Kruskal–Wallis test followed by Dunn’s multiple comparisons test was applied. To assess differences in cell proportions or categorical distributions across experimental conditions, Chi-square (χ²) tests were used. When applicable, exact *P*-values were calculated and reported. In figures, statistical significance is indicated as: P < 0.05 (*), P < 0.01 (**), P < 0.001 (***), and P < 0.0001 (****).

## Data availability

Single-cell RNA-seq data have been deposited in the NCBI Sequence Read Archive (SRA) under the BioProject accession PRJNA1330728 (https://www.ncbi.nlm.nih.gov/bioproject/). Primary source data and original images are available on Figshare, as detailed in Supplementary table 4. All other data supporting the findings of these studies are included in the manuscript as supplementary dataset and are also available from the corresponding author upon reasonable request.

## Supplementary data

### Supplementary figures

**Supplementary figure 1.**
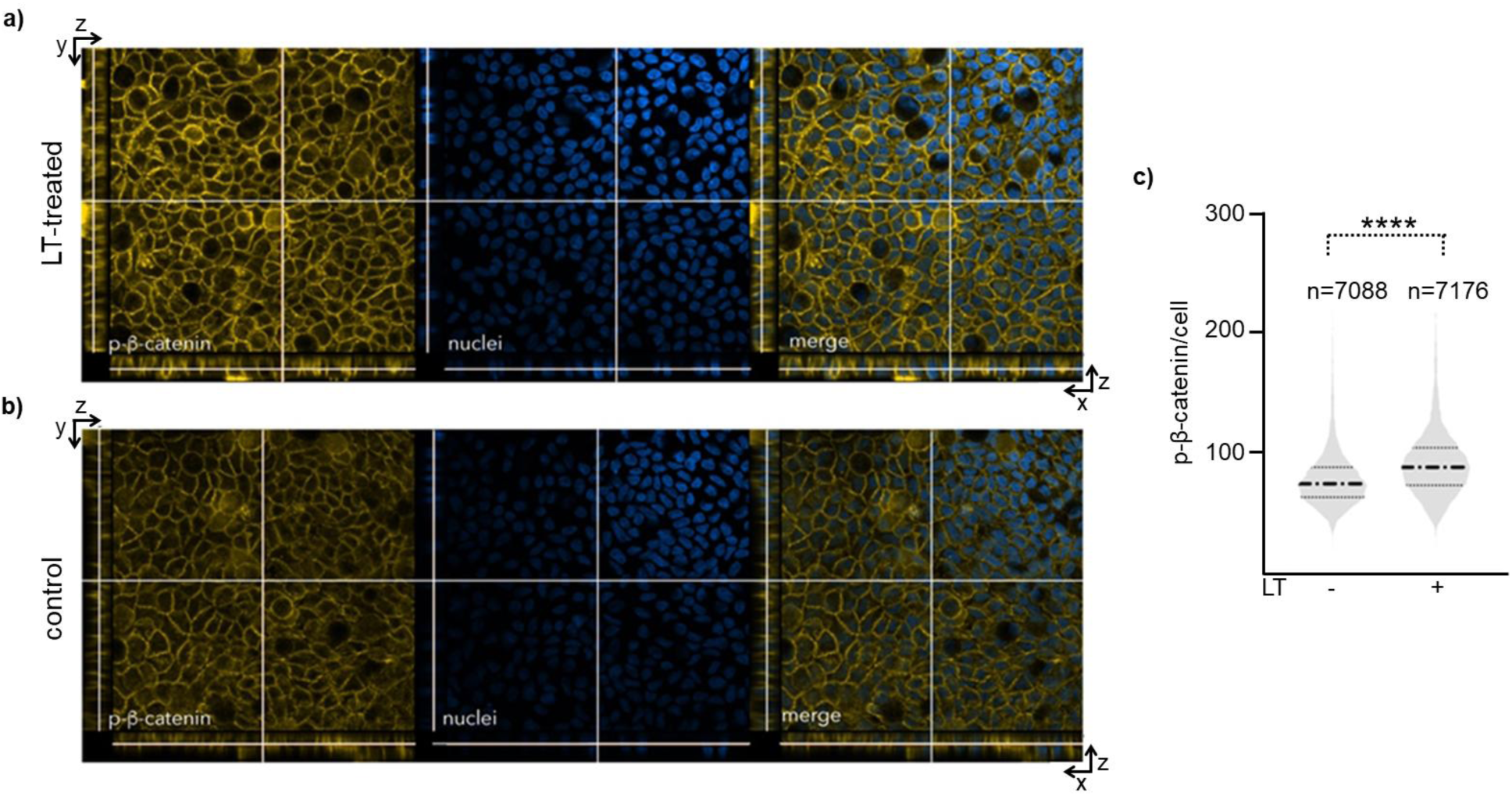
Phophorylated β-catenin (Ser_675_) accumulates in target intestinal cells following treatment with heat-labile toxin. **a.** Top panels show phospho-β-catenin accumulation in representative field of LT-treated Hu235D cells compared to untreated cells (**b**). **c.** Violin plots at right summarize p-β-catenin Ser_675_ signal per cell obtained from n=7088 cells in control, and n=7176 cells in LT-treated group. Data was accumulated over 20 randomly selected fields. Heavy dashed lines in each group represent median values, thin lines are quartiles. ****p<0.0001 (Mann-Whitney).

**Supplementary figure 2.**
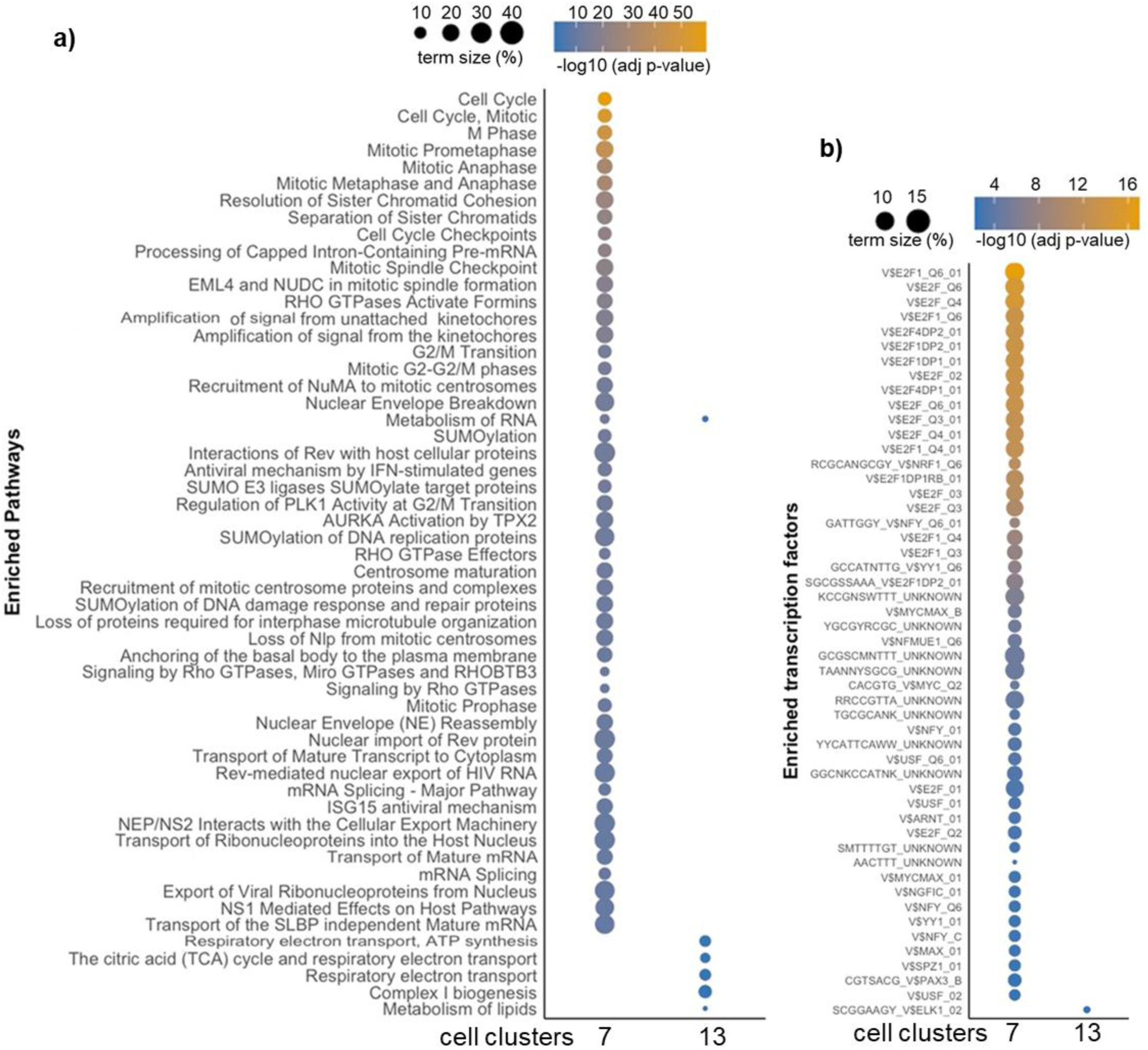
Over-representation analysis (ORA) of cluster 7 (TA) and cluster 13 marker genes (BEST4+EC). Dot plots displaying significantly enriched (FDR < 0.05) pathways (a) and transcription factors (b) along the Y-axis, among cluster marker genes for clusters 7 and 13. Dot size indicates the number of marker genes contributing to each pathway, and color reflects the adjusted statistical significance (−log10 FDR), with yellow indicating stronger enrichment and blue indicating lower significance.

**Supplementary figure 3.**
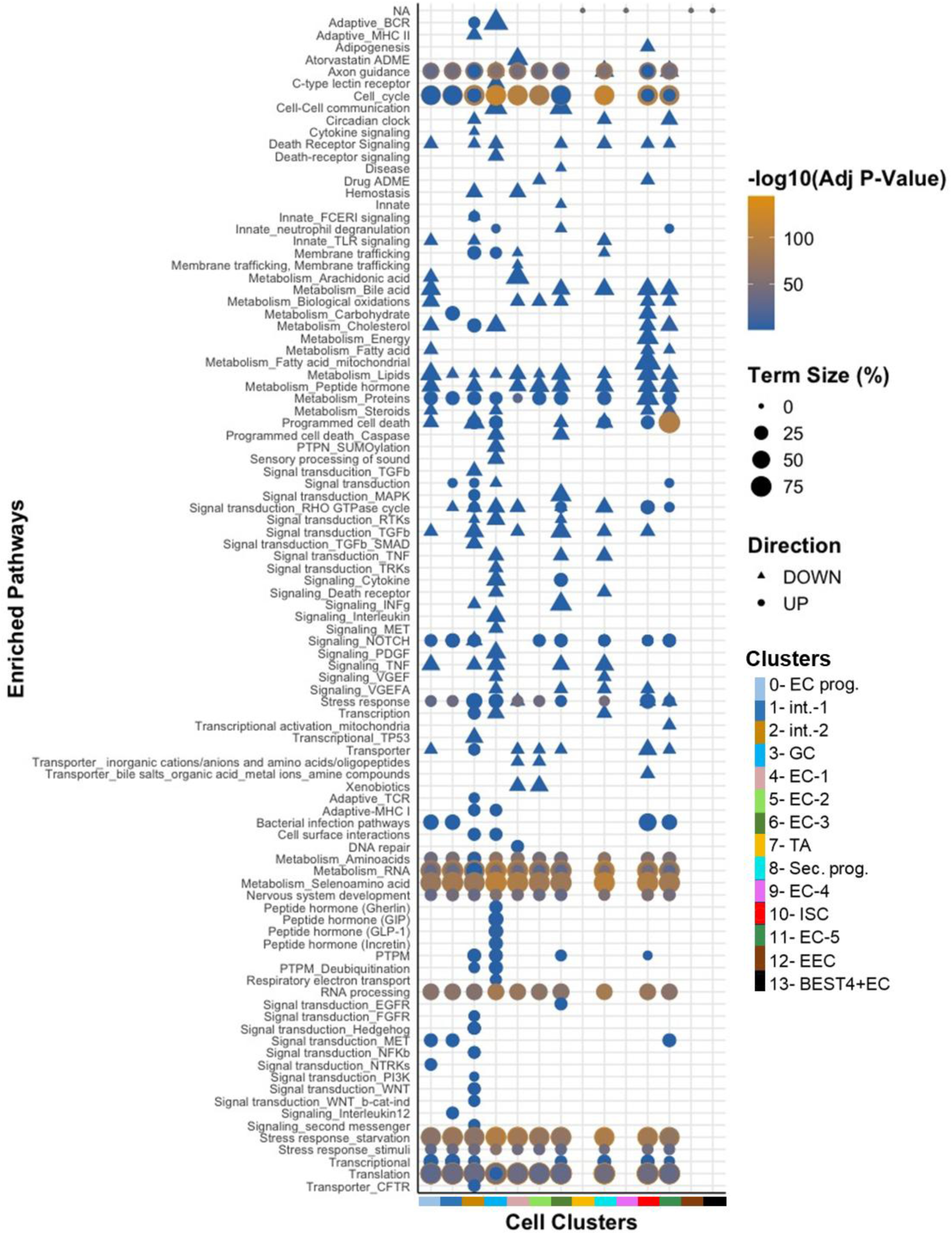
Over-representation analysis (ORA) of significantly up- and down-regulated genes between LT and control samples. The dot plot displays significantly enriched biological pathways (Y-axis) across 14 epithelial cell clusters (X-axis) among differentially expressed genes in LT-treated vs. control intestinal organoids (FDR <0.05). Circles represent upregulated pathways, while triangles represent downregulated pathways. Dot size indicates the number of DEGs contributing to each pathway, and color reflects the adjusted statistical significance (−log10 FDR), with yellow indicating stronger enrichment and blue indicating lower significance.

**Supplementary figure 4.**
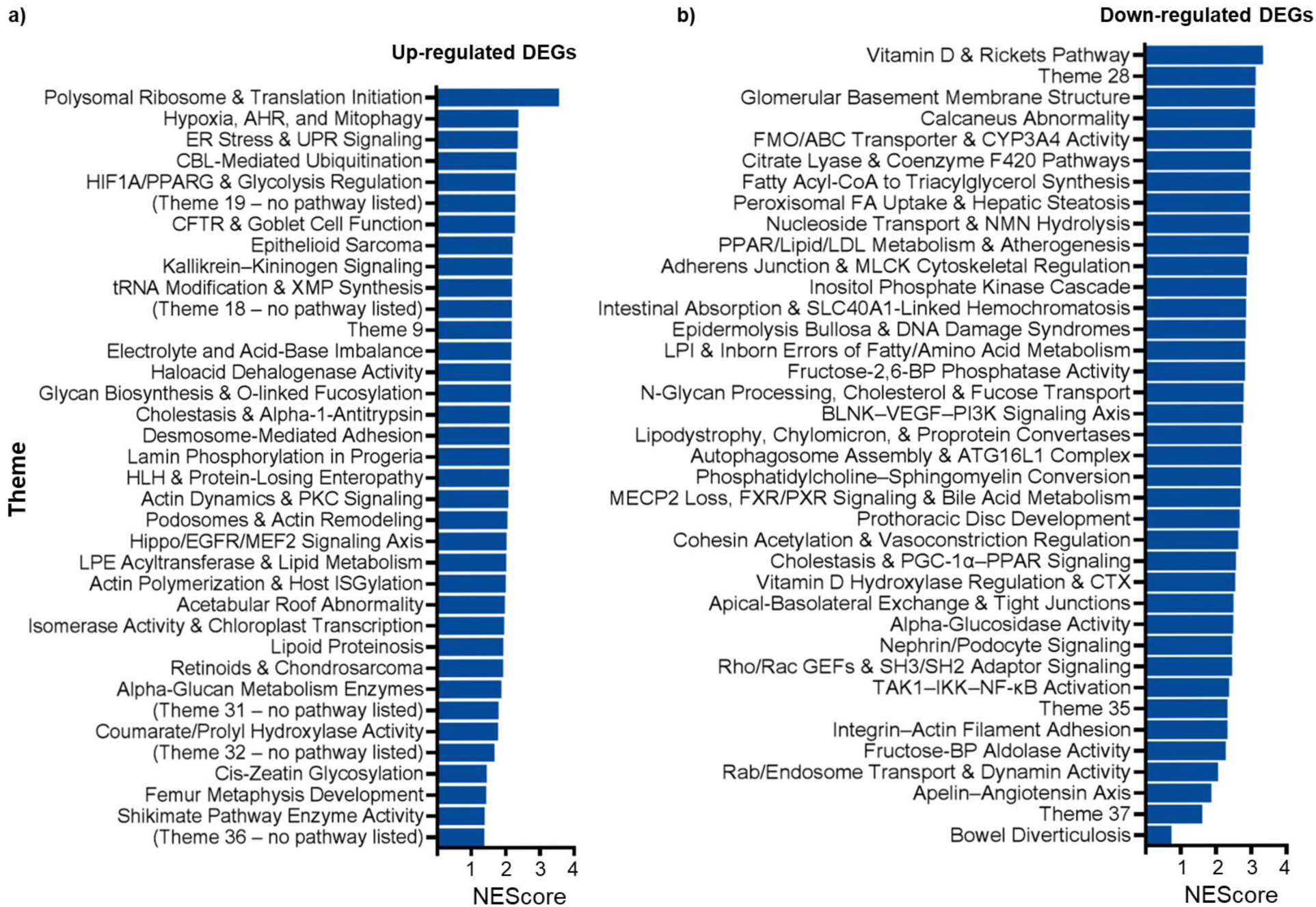
CompBio analysis of differentially expressed genes (DEGs) in LT-treated enteroids. a) Bar plot showing themes significantly enriched among downregulated genes (FDR-adjusted p < 0.05). b) Bar plot showing themes significantly enriched among upregulated genes (FDR-adjusted p < 0.05). Pathway enrichment was conducted using CompBio v2.9 with default parameters. NEScore represents the normalized enrichment score derived from CompBio theme association analysis. Themes were prioritized based on total score. See Supplementary dataset for full theme descriptions and gene lists.

**Supplementary figure 5.**
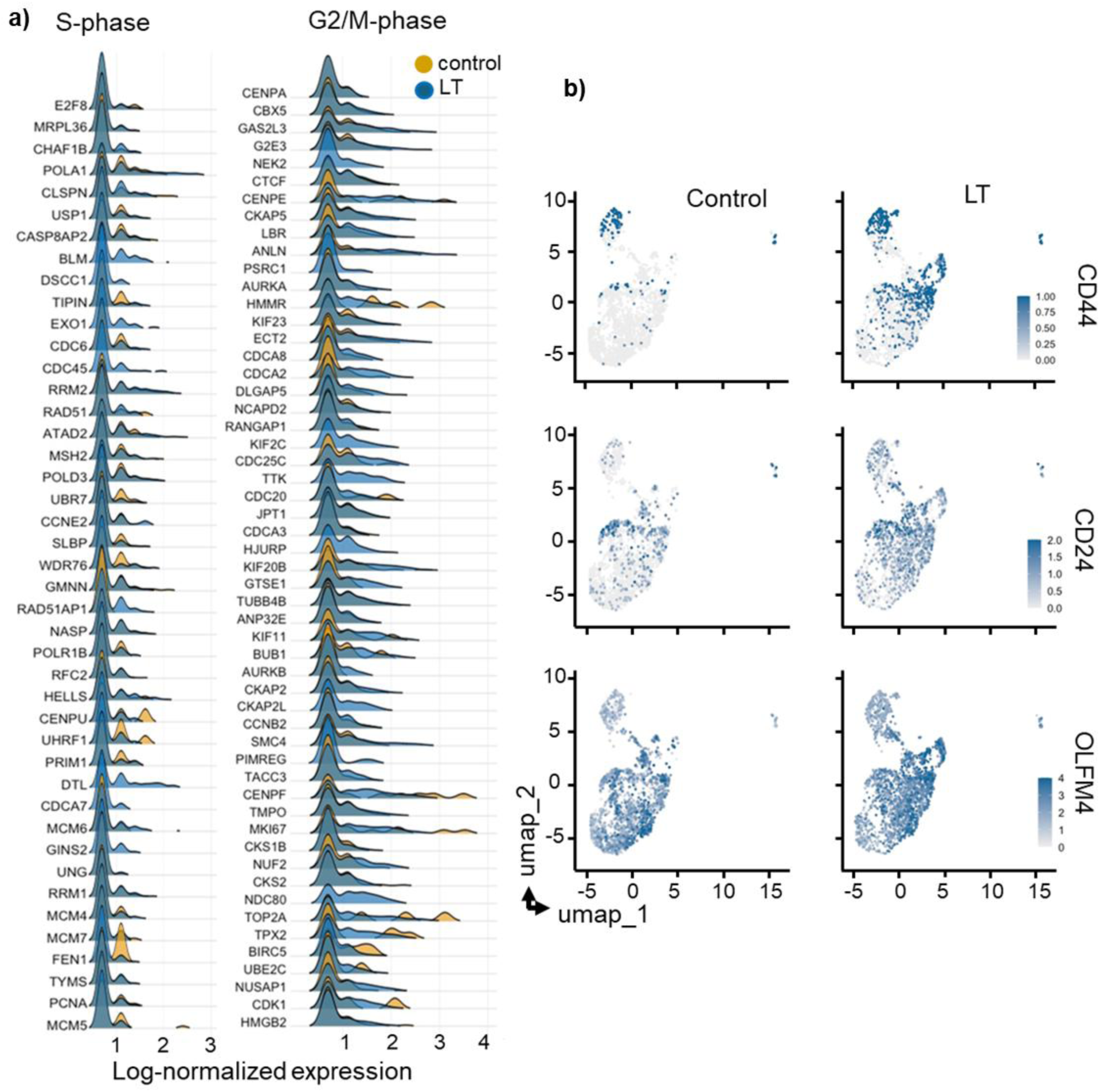
Expression of cell-cycle phase and stem/progenitor genes. a) Ridge plots showing expression of cell cycle phase marker genes. Normalized expression of S-phase genes (left) and G2/M-phase genes (right) in control and LT-treated enteroids. b) Feature plots displaying expression of CD44, CD24, and OLFM4 markers of stem or undifferentiated states across cell clusters in control (left) and LT-treated (right) enteroids.

**Supplementary figure 6.**
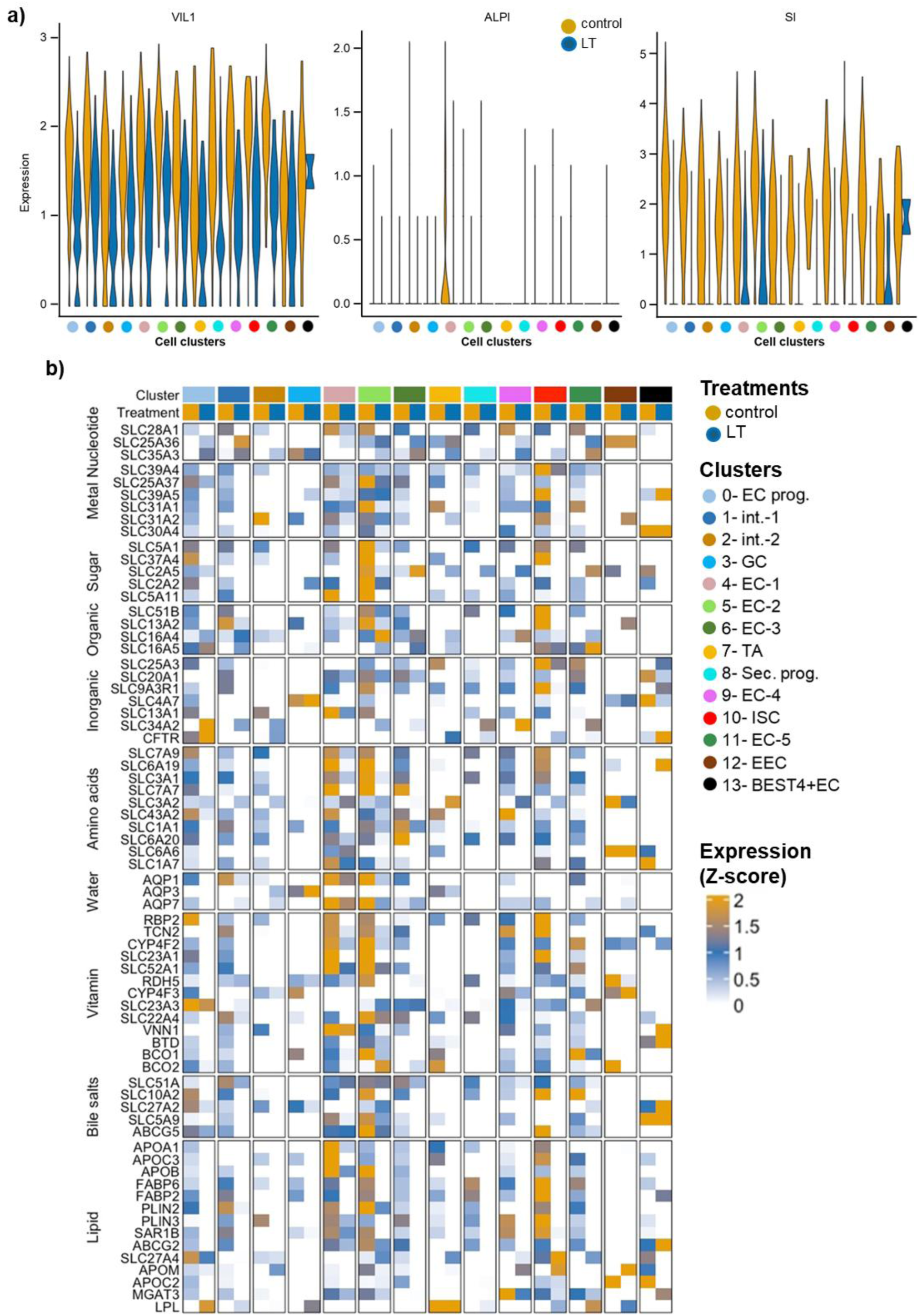
LT impairs enterocyte maturation and suppresses expression of nutrient transporters. **a)** Violin plots showing gene expression profile of mature enterocyte markers (VIL1, ALPI, and SI) in control and LT-treated human ileal enteroids. Each panel displays normalized expression levels across the epithelial cell clusters. **b)** Heatmap displaying expression profiles of nutrient transporter genes across all clusters in control and LT-treated human ileal enteroids. Expression values represent Z-score of SCTransform-normalized data averaged per cluster. Bars above the heatmap denote treatment (control, LT) and cluster identities as indicated.

**Supplementary figure 7.**
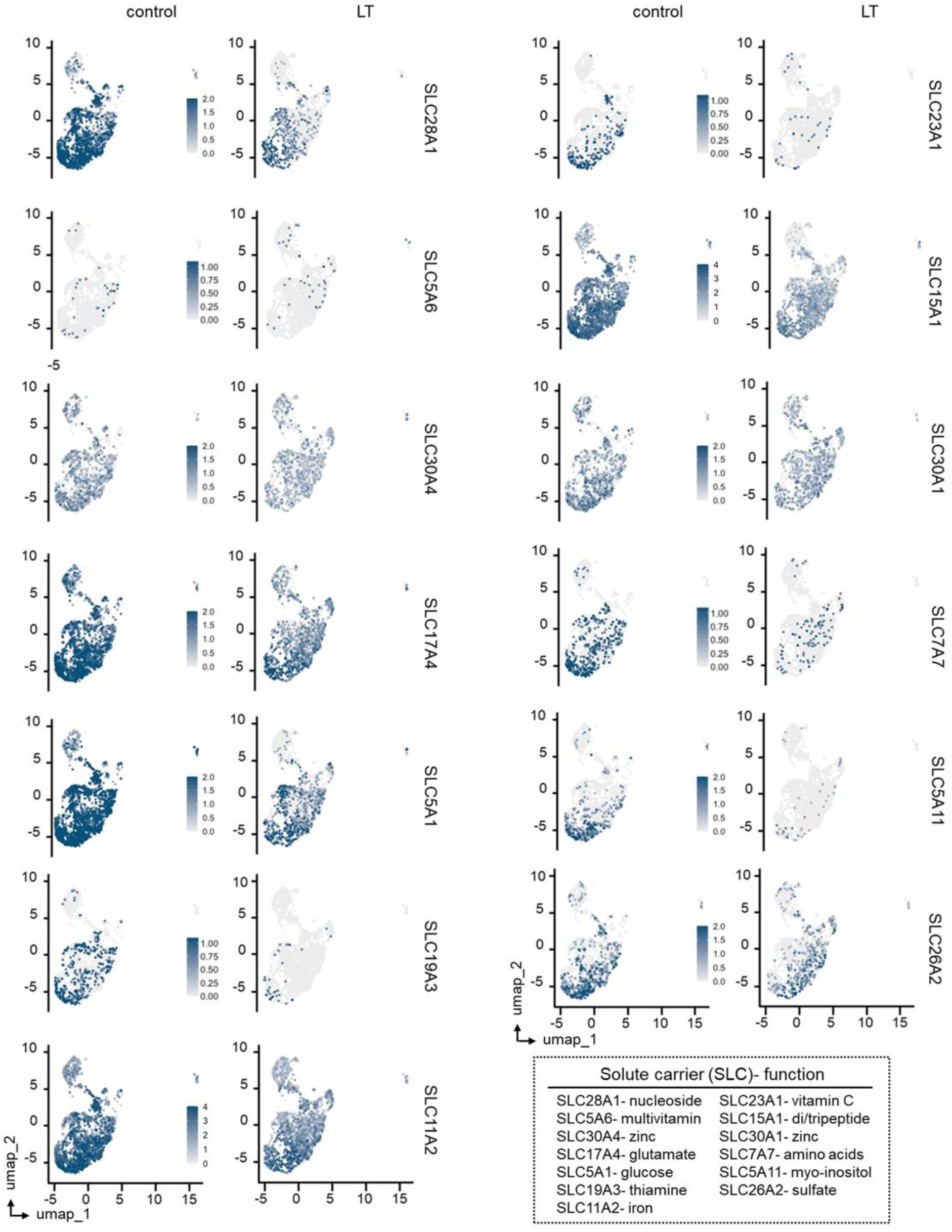
LT suppresses expression of multiple nutrient transporter genes in human ileal enteroids. UMAP feature plots showing the expression of select solute carrier (SLC) family genes involved in nutrient absorption across control (left panels) and LT-treated (right panels) human ileal enteroids. Each pair of panels displays normalized expression levels of a specific SLC gene projected onto the epithelial cell landscape. Gene function annotations are summarized in the accompanying table.

### Supplementary tables

**Supplementary table 1:**
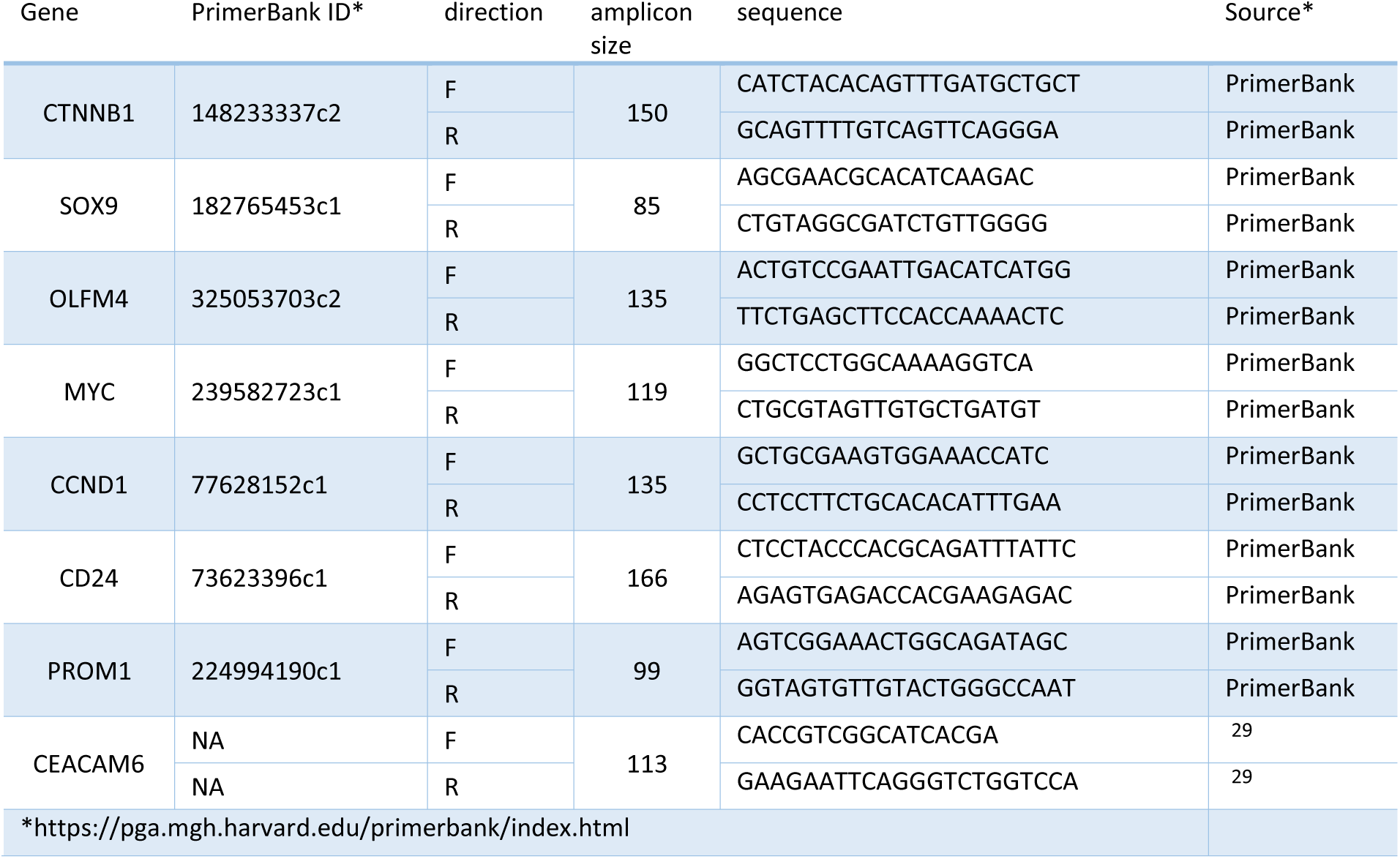
Primer sequences used in Quantitative RT-PCR experiments.

**Supplementary table 2:** Antibody descriptions and sources for immunoblot and fluorescence microscopy experiments.

| Protein, antigen | clone<sup>1</sup> | host | isotype | conjugation | source | RRID* |
| --- | --- | --- | --- | --- | --- | --- |
| GAPDH, C terminal peptide | m \| 14C10 | Rabbit | IgG |  | Cell Signaling 2118S | AB 561053 |
| GSK-3a/b (synthetic peptide surrounding Gln269 of human GSK-3a protein) | D75D3 | Rabbit | IgG |  | Cell Signaling 5676 | AB 10547140 |
| GSK-3b Ser<sub>9</sub> phospho-peptide | polyclonal | Rabbit | IgG |  | Cell Signaling 9336S | AB 331405 |
| GSK-3a/b (Ser21/9) synthetic phosphopeptides | Polyclonal | Rabbit |  |  | Cell Signaling 9331 | AB 329830 |
| b-catenin (synthetic peptide circa Pro174 of human protein) | monoclonal (D10A8) | Rabbit | IgG |  | Cell Signaling 8480 | AB 11127855 |
| b-catenin Ser<sub>675</sub> C-terminal phospho-peptide | polyclonal | Rabbit | IgG |  | Invitrogen PA5-77933 | AB 2736355 |
| MKI67 | polyclonal | Rabbit | IgG |  | Novus NB500-170 | AB 10001977 |
| Villin (BDID2C3) | monoclonal | Mouse | IgG1 |  | Santa Cruz sc-66022 | AB 2216247 |
| ALPI | polyclonal | Rabbit | IgG |  | Invitrogen PA5-22210 | AB 11153471 |
| GAPDH (14C10) | monoclonal | Rabbit |  |  | Cell Signaling 2118S | AB 561053 |
| Lamin B1 | monoclonal (EPR22165-121) | Rabbit | IgG |  | Abcam ab229025 | AB 3083735 |
| Anti-rabbit IgG | polyclonal | Goat | IgG (H+L) | Alexa Fluor 488 | ThermoFisher A-11070 | AB 143165 |
| Anti-rabbit IgG | polyclonal | Goat | IgG | HRP | Cell Signaling 7074S | AB 2099233 |
| Anti-mouse IgG | polyclonal | Horse | IgG | HRP | Cell Signaling 7076S | AB 330924 |

\* https://rrid.site/

**Supplementary table 3:** Plasmids used in the construction of TCF/LEF reporter enteroid lines.

| plasmid | description | source | reference |
| --- | --- | --- | --- |
| psPAX2 | 2<sup>nd</sup> generation lentiviral packaging plasmid | addgene<sup>1</sup> |  |
| pMD2.G | VSV-G envelope expressing plasmid for lentiviral packaging | addgene<sup>2</sup> |  |
| p7TFP | 7X copies of the TCF/LEF transcriptional response element GATCAAAGG linked to firefly luciferase ; puromycin R | addgene<sup>3</sup> | <sup>30</sup> |

<sup>1</sup>psPAX2 was a gift from Didier Trono (https://www.addgene.org/12260/; RRID:Addgene\_12260)
<sup>2</sup>pMD2.G was a gift from Didier Trono (https://www.addgene.org/12259/ ; RRID:Addgene\_12259)
<sup>3</sup>p7TFP was a gift from Roel Nusse (https://www.addgene.org/24308/; RRID:Addgene\_24308)

**Supplementary table 4:** links to the original images and datasets.

| Type | Description | DOI |
| --- | --- | --- |
| Figure 2b-c. | Original blots of β-catenin, phospho-β-catenin- Ser<sub>675</sub>, GSK3β, phospho-GSK3β-Ser<sub>9</sub>, Lamin-b1, and GAPDH | 10.6084/m9.figshare.30100273 |
| Figure 6a. | Original blot of intestinal alkaline phosphatase (ALPI) | 10.6084/m9.figshare.30104512 |
| Figure 6b. | Original blot of Villin | 10.6084/m9.figshare.30104635 |
| Data | Data associated with graphs | 10.6084/m9.figshare.30198553 |
| Data | Data associated with tables | 10.6084/m9.figshare.30201280 |

## Acknowledgements

JMF was supported by funding from the Department of Veterans Affairs (VA) (5I01BX001469-05), and the National Institute of Allergy and Infectious Diseases (NIAID) of the National Institutes of Health (NIH) R01 AI170949, R01 AI089894, and support by the NIH Washington University DDRCC Grant NIDDK P30 DK052574. AS was supported by National Institute of Allergy and Infectious Diseases of the National Institutes of Health under Award Number T32AI007172 and JIT1052 (supported by the Washington University Institute of Clinical and Translational Sciences grant UL1TR002345 from the National Center for Advancing Translational Sciences (NCATS) of the NIH. We thank the Genome Technology Access Center at the McDonnell Genome Institute at Washington University School of Medicine for help with genomic analysis. The Center is partially supported by NCI Cancer Center Support Grant #P30 CA91842 to the Siteman Cancer Center from the National Center for Research Resources (NCRR), a component of the NIH, and NIH Roadmap for Medical Research. This publication is solely the responsibility of the authors and does not necessarily represent the official view of NCRR, NIH, or VA.

